# Bioinformatic inference of the exercise-responsive control of p70 S6 kinase through *RPS6KB1* expression

**DOI:** 10.1101/2025.10.11.681841

**Authors:** Taylor J. McColl, Renfei Zhang, Aurelien Dugourd, Julio Saez-Rodriguez, David C. Clarke

## Abstract

Previous research suggests that the absolute levels of p70 S6 kinase (p70S6K) are a key determinant of the rate of skeletal muscle protein synthesis (MPS). p70S6K levels are in part determined by the transcriptional control of the gene encoding p70S6K, *RPS6KB1*, but the molecular mechanisms governing its expression are poorly understood. The purpose of this study was to infer the molecular regulatory network governing *RPS6KB1* expression. We applied a novel bioinformatic network inference algorithm called CARNIVAL (CAusal Reasoning pipeline for Network identification using Integer VALue programming) to infer the signaling network downstream of canonical exercise sensors controlling *RPS6KB1*-specific transcription factors (TFs) after acute aerobic (AE) or resistance exercise (RE). CARNIVAL integrates a prior knowledge network, TF and signaling pathway activities inferred from transcriptomic data, and perturbation targets to predict the network that best explains the data. The networks revealed intracellular sensors and hormone receptors controlling *RPS6KB1*-specific TFs. Both exercise types resulted in AMPK-mediated *SNAI1* regulation, but *HIF1A* was distinctly controlled (AE: PHD1-3, FIH; RE: AMPK). AE controlled *FOXA1* via insulin, TGF-β, and myostatin signalling, while RE controlled *CEBPA* via MAP3Ks. Our study is the first to apply a comprehensive bioinformatic network inference algorithm to infer causal exercise-responsive signaling networks. The results of our analysis motivate experimentally testable hypotheses pertaining to the molecular control of *RPS6KB1* transcription in human skeletal muscle in response to aerobic and resistance exercise.

## Introduction

Skeletal muscle is essential for health and quality of life (Kraemer & Ratamess, 2004; Karagounis & Hawley, 2010). Muscle function is in part tied to its mass (Cormie *et al*., 2011). Maintaining or increasing muscle mass benefits health, whereas loss of muscle mass can increase rates of morbidity and mortality (McLeod *et al*., 2016). Loss of muscle mass occurs in disease states such as sarcopenia, which is the age-related loss of muscle mass and function (Kirk *et al*., 2024), Those with sarcopenia are hospitalized more frequently (Beaudart *et al*., 2017) and impose a significant financial burden on public healthcare (Janssen *et al*., 2004; Sousa *et al*., 2016; Steffl *et al*., 2017). Consequently, improved understanding of skeletal muscle mass regulation and effective therapies to counter the age-related loss of muscle mass are needed.

Skeletal muscle mass is primarily composed of myofibrillar proteins (e.g., actin, myosin) (Roberts *et al*., 2020), such that its mass is dependent on its protein content. Muscle proteins are continuously degraded and synthesized (Atherton & Smith, 2012), with their levels determined by the balance between the rates of muscle protein synthesis (MPS) and muscle protein breakdown (MPB). Of these, MPB is relatively constant and MPS is responsive to anabolic stimuli (Phillips *et al*., 2005; Drummond & Rasmussen, 2008; Morton *et al*., 2015; Deutz *et al*., 2025). Consequently, muscle protein balance, i.e. the difference between MPS and MPB, is primarily dependent on changes to MPS.

Muscle protein balance is controlled by several stimuli that modulate MPS and MPB, (Hoppeler, 2016). Anabolic hormones, such as insulin, growth hormone, insulin-like growth factor 1, and testosterone, positively control MPS and promote muscle protein balance (Rooyackers & Nair, 1997; Hoppeler, 2016). Furthermore, insulin increases muscle protein balance via anticatabolic mechanisms that reduce the rates of MPB (Rabinowitz & White, 2010; Neel *et al*., 2013; McKendry *et al*., 2021). Exercise promotes muscle protein balance through mechanical loading across the muscle (Rindom & Vissing, 2016). Appropriately programmed resistance exercise is undeniably the most potent nonpharmacological stimulator of skeletal muscle hypertrophy (Lim *et al*., 2022). Aerobic exercise can likewise promote muscle hypertrophy, albeit less potently than resistance exercise (Konopka & Harber, 2014; Grgic *et al*., 2019). Each stimulus transiently modifies the rates of MPS and MPB for up to 24 hours (Burd *et al*., 2011; Trommelen *et al*., 2023), such that they need to be applied at consistent intervals to maintain or increase muscle protein balance.

Signaling mediated by anabolic stimuli predominantly converges on the mechanistic target of rapamycin complex 1 (mTORC1) (Hoppeler, 2016). mTORC1 is a protein kinase that integrates anabolic signaling responses and modulates MPS by activating downstream targets such as p70 S6 kinase (p70S6K) (Ma & Blenis, 2009; Magnuson *et al*., 2012; Wu *et al*., 2022). p70S6K controls translation initiation, which is considered the “rate-limiting step” for protein biosynthesis (Ma & Blenis, 2009). mTORC1-mediated phosphorylation and activation of p70S6K liberates eIF3, enabling pre-initiation complex assembly (Kleijn *et al*., 1998; Ma & Blenis, 2009; Magnuson *et al*., 2012). Activated p70S6K also controls various downstream physiological and pathological processes, including cell size, metabolism, proliferation, and survival (Wu *et al*., 2022).

The critical role of p70S6K is evident in knockout studies, wherein its loss leads to muscle atrophy (Ohanna *et al*., 2005; Aguilar *et al*., 2007), impaired force-generating capacity and muscle structure (Marabita *et al*., 2016), and blunted responses to hypertrophic stimuli (Ohanna *et al*., 2005). Furthermore, total p70S6K protein concentrations in older men were found to be 50% of those in young men, corresponding to reduced MPS rates following amino acid ingestion, an anabolic stimulus sensed by mTORC1 (Cuthbertson *et al*., 2005). Computational modeling analyses of human skeletal muscle have also identified total p70S6K levels as important for controlling MPS (McColl & Clarke, 2024; McColl *et al*., 2025). Therefore, understanding how p70S6K levels are controlled in skeletal muscle is critical for health and muscle function.

While the post-translational control of p70S6K is well-documented, particularly in terms of the phosphorylation of key residues (Ma & Blenis, 2009; Magnuson *et al*., 2012; Wu *et al*., 2022), the molecular mechanisms underlying its transcriptional control remain incompletely understood. The *RPS6KB1* gene, located on chromosome 17 (17q23), encodes two S6K proteins, p70S6K and p85S6K, through alternative translation start sites (Wu *et al*., 2022; Perez *et al*., 2025). *RPS6KB1* expression is transcriptionally controlled by the binding of transcription factors (TFs) to specific promoter and enhancer regions of the DNA.

TF-mediated control of *RPS6KB1* has been investigated in breast and brain cancers. In estrogen receptor-positive breast cancer cells, estrogen activates *RPS6KB1* expression via estrogen receptor α (ERα) (Maruani *et al*., 2012). ERα interacts with the *RPS6KB1* promoter region with the involvement of the transcriptional factor GATA3 (Maruani *et al*., 2012). Conversely, the estrogen-related receptor alpha (ERRα) negatively regulates *RPS6KB1* expression in triple-negative breast cancer cells by binding to its promoter region (Berman *et al*., 2017). Additionally, evidence suggests that *RPS6KB1* expression in brain tumors is controlled by hypoxia-induced genes (Ismail, 2012). Despite these insights, the full range of molecular mechanisms governing *RPS6KB1* transcription remains poorly understood. Given the complexity of transcriptional control, systematically characterizing individual TFs and their control of target genes is both tedious and time-consuming, highlighting the need for a more comprehensive and efficient global approach.

Bioinformatic methods have been developed to investigate the transcriptional control of key genes. In exercise biology, applications have largely focused on data-driven, descriptive transcriptomics, including differential expression and enrichment analyses (Keller *et al*., 2011; Dickinson *et al*., 2018), pathway-level inference (Hoffman *et al*., 2015; Robinson *et al*., 2017), network-based methods (Phillips *et al*., 2013; Lindholm *et al*., 2014), and meta-analysis approaches (Pillon *et al*., 2020, 2022). To the best of our knowledge, no approaches in exercise biology have incorporated causal mechanistic information to infer the upstream signaling events that drive the observed transcriptional responses. CARNIVAL (Causal Reasoning pipeline for Network identification using Integer VALue programming) (Liu *et al*., 2019) is a bioinformatic network inference algorithm that integrates prior knowledge of signaling networks with transcriptomic data through causal reasoning to generate mechanistic hypotheses of upstream events controlling observed transcriptional responses. CARNIVAL and its derivatives promise to serve as unique and powerful tools for studying exercise biology because they can identify potential intervention targets by mapping how upstream signals propagate to downstream transcriptional responses.

The purpose of this study was to explore how *RPS6KB1* is controlled by transcription factors following aerobic and resistance exercise by applying a bioinformatic network inference approach to publicly available transcriptomic data. We identified TFs controlling *RPS6KB1* expression using a curated prior knowledge network (Türei *et al*., 2016, 2021). Using these *RPS6KB1*-specific TFs, we applied the network inference tool CARNIVAL to infer the molecular regulatory network downstream of canonical exercise sensors that controls these *RPS6KB1*-specific TFs (Liu *et al*., 2019).

## Materials and Methods

### Approach to the problem

The network inference approach involved three general steps. First, we identified TFs controlling *RPS6KB1* using curated databases. We then inferred the activities of these TFs from publicly available exercise transcriptomic datasets. Next, we applied CARNIVAL, which integrates inferred TF activities with prior knowledge of canonical signaling networks to predict the upstream exercise sensors controlling these TFs. This approach enabled the identification of key exercise sensors controlling TFs that target *RPS6KB1*, as well as the signaling pathways through which they operate, thereby highlighting potential pharmacological or interventional targets for modulating *RPS6KB1* expression.

### Identification of transcription factors controlling *RPS6KB1*

We used the OmniPath (Türei *et al*., 2021) curated signaling network to identify TFs controlling *RPS6KB1*. Specifically, we used the Cytoscape OmniPath App (Shannon *et al*., 2003; Ceccarelli *et al*., 2020) to generate a comprehensive network of human TF-target interactions, with TF confidence scores ranging from A to E. We identified the *RPS6KB1* node in the network and selected its “first neighbors” to map the up- and downstream signaling network. Any immediate upstream neighbors of *RPS6KB1* were considered TFs that directly control *RPS6KB1* expression.

### Transcriptomic data

We used the transcriptomic dataset from Dickinson et al. (Dickinson *et al*., 2018) as an input to the bioinformatic network inference tool CARNIVAL (Liu *et al*., 2019). The study measured the transcriptomic response in human skeletal muscle following acute aerobic and resistance exercise using a randomized crossover design (Dickinson *et al*., 2018). Six healthy, recreationally active men (aged 27±3 years) completed both a resistance exercise bout (isotonic unilateral leg extension: 8 sets of 10 repetitions at 60-65% of one-repetition maximum, 3-minute inter-set rest) and an aerobic exercise bout (40 minutes of stationary cycling at 70% of maximal heart rate), with a 9±3-day washout period between sessions (Dickinson *et al*., 2018). Muscle biopsies were collected before the first exercise bout and at 1 and 4 hours post-exercise for each session (Dickinson *et al*., 2018). Whole transcriptome RNA sequencing was then performed using Illumina HiSeq 2500 (Dickinson *et al*., 2018).

### Data preprocessing

The dataset from Dickinson et al. was provided as individual .txt files for each biopsy time points for each participant, containing transcript read counts for each gene. The data were imported into R and tabulated by time point and participant. We converted all 0 counts to “NA” and then applied log2 transformation. The transformed data were normalized using the “vsn” package in R (Huber *et al*., 2002), which applies variance-stabilizing normalization to standardize variance across signal intensities.

### Differential gene expression analysis

We identified differentially expressed genes using the R package “limma” (Ritchie *et al*., 2015) across five conditions: baseline, 1 hour post-aerobic exercise (AT.1h), 1 hour post-resistance exercise (RT.1h), 4 hours post-aerobic exercise (AT.4h), and 4 hours post-resistance exercise (RT.4h). Specifically, we fit four linear models to evaluate the differential gene expression between baseline and each time point (AT.1h, AT.4h, RT.1h, and RT.4h). For each comparison, we obtained t-values and p-values, adjusting the latter to control the false discovery rate (FDR). Finally, we created quantile-quantile plots to assess the gene expression magnitudes across the four conditions relative to baseline.

### Inference of transcription factor activities

Transcription factor activity was estimated from the gene expression of their target genes using Discriminant Regulon Expression Analysis (DoRothEA) (Garcia-Alonso *et al*., 2019). DoRothEA is a curated collection of TFs and their transcriptional targets, known as regulons, derived from various sources of evidence (e.g., literature, ChIP-seq data, TF binding site predictions, and gene expression inference) (Garcia-Alonso *et al*., 2019). Each TF-target interaction in DoRothEA is assigned a confidence score ranging from A (highest confidence) to E (lowest confidence), based on the amount and reliability of supporting evidence (Garcia-Alonso *et al*., 2019). To ensure a comprehensive set of candidate TFs potentially controlling *RPS6KB1* transcription, we retained all regulons with confidence scores from A to E, resulting in 1,333 unique TFs with 454,505 TF-target interactions. Since the inferred TF activities are based on the expression of target genes, we applied an additional filtering step, retaining only TFs with at least three downstream targets (i.e., TF-target interactions). This ensured that inferred TF activities were derived from multiple gene targets rather than being disproportionately influenced by a single or two TF-target interactions. Importantly, this filtering process retained all *RPS6KB1*-specific TFs.

Next, we inputted the differential expression t-values derived from the “limma” package, along with the filtered regulon subset, into the “run_viper” function of the Virtual Inference of Protein-activity by Enrichment Regulon analysis (VIPER) (Alvarez *et al*., 2016) R package. The “run_viper” function calculates the normalized enrichment scores (NES) for each TF based on these inputs. The NES infers the activity of the TF by the collective expression changes of its target genes, reflecting its regulatory effect.

### Inference of signaling pathway activities

We then used the Pathway RespOnsive GENes for activity inference (PROGENy) (Schubert *et al*., 2018) tool to infer signaling pathway activities that may serve as drivers of the observed transcriptomic response. PROGENy estimates pathway activities for 14 canonical pathways commonly evaluated in perturbation experiments, such as NF-κB, TGF-β, PI3K, MAPK, and others (Schubert *et al*., 2018). We filtered the differential expression t-values from “limma” to retain the top 100 most responsive genes and inputted those values into PROGENy to derive pathway scores. Each pathway score is calculated using predefined sets of pathway-responsive genes, applying a linear weighting model to their expression levels to infer the activity of specific signaling pathways in the sample. The resulting pathway scores are normalized to a mean of zero and a standard deviation of one.

### Inference of the molecular regulatory network

CARNIVAL uses causal reasoning to integrate TF activities with upstream regulatory networks to predict the network that best explains the data (Liu *et al*., 2019). CARNIVAL requires as inputs a prior knowledge network (PKN) and transcription factor activities inferred from transcriptomic data (e.g., DoRothEA) (Liu *et al*., 2019). Optional inputs include signaling pathway activities (e.g., PROGENy) and a list of *perturbation targets*, which are proteins, pathways, or regulators expected to have altered activities in response to experimental treatments or environmental conditions (Liu *et al*., 2019).

The CARNIVAL objective function includes *alpha weight* and *beta weight* parameters: alpha weight controls how closely node (i.e., gene) activities follow prior knowledge (i.e., the difference between the inferred and prior knowledge inferred node activities), while beta weight controls network sparsity by controlling the number of edges included in the network (Liu *et al*., 2019). We set alpha weight and beta weight to their default values of 1 and 0.2, respectively (Liu *et al*., 2019).

We used the signed (i.e., stimulatory or inhibitory) and directed (i.e., indicating the direction of influence between molecules) human signaling network from OmniPath (Türei *et al*., 2016) as the PKN. This network contains 9,306 signed and directed edges connecting 3,610 proteins, which collectively represent human regulatory and signaling pathways. We imported the OmniPath PKN on February 7^th^, 2025, using the “import_omnipath_interactions” function with default parameters from the “OmnipathR” R package. The TF activities for the *RPS6KB1*-specific TFs from DoRothEA and the signaling pathway activity scores inferred from PROGENy were also used as inputs for CARNIVAL.

CARNIVAL can be run as two distinct pipelines: 1) Inverse CARNIVAL (invCARNIVAL), which does not require information about perturbation targets, and 2) Standard CARNIVAL (stdCARNIVAL), which incorporates known perturbation targets to guide network inference. We used stdCARNIVAL to generate inferred networks linking *RPS6KB1*-specific TFs to canonical exercise sensors, ensuring relevance to exercise-mediated mechanisms. Specifically, we provided CARNIVAL a comprehensive list of 50 canonical exercise sensors as perturbation targets (Supplementary Table 1). This list included intracellular sensor proteins such as AMPK isoforms, CaMKs, MAP3Ks, mechanosensors, and prolyl hydroxylases, and hormonal sensors, including androgen receptor, insulin receptor, and TGF-β receptors. Designating these exercise sensors as perturbation targets ensured that the inferred signaling network would extend from *RPS6KB1*-specific TFs to these upstream exercise sensors, providing insight into how exercise regulates *RPS6KB1* transcriptional activity. Additionally, we applied invCARNIVAL to identify possible non-canonical upstream regulators that may influence the exercise-mediated control of *RPS6K1*-specific TFs beyond the canonical exercise sensors.

CARNIVAL infers the signaling network topology and node activities by formulating the problem as an optimization task using integer linear programming (ILP). We used the ILP solver, CPLEX (complex linear programming expert), to integrate the predicted TF and pathway activity scores with the PKN to identify the optimal subnetwork. This subnetwork is a directed and signed subset of the PKN that best aligns the experimental data while satisfying causal constraints, which encode activation or inhibition relationships. By linking the TF and pathway activities to upstream targets of perturbation, this approach minimizes inconsistencies between the inferred network and the experimental data, resulting in a biologically plausible regulatory model.

The CPLEX optimizer identifies multiple high-quality networks, rather than a single optimal solution. Using an aggressive search strategy, CPLEX generates a diverse pool of solutions, which is then narrowed down to the top 100 most diverse subnetworks. As a result, the final CARNIVAL-inferred network integrates these diverse, high-quality subnetworks, typically combining 100 of them to form the final network. The consistency of the network pool is summarized by the *edge weights* and the *node attribute values* in the final network, which are determined as follows:

- Edge weights: percentage of subnetworks containing a specific interaction
- Node attribute values:

`o` zeroAct: percentage of subnetworks where the node has zero activity (i.e., not included in the network solution)
`o` upAct: percentage of subnetworks where the node has positive activity
`o` downAct: percentage of subnetworks where the node has negative activity
`o` avgAct: the difference between upAct and downAct, representing the average activity.
`o` nodeType: nodes identified as perturbation targets are tagged with a “S”, while measured TFs are tagged with a “T”.

### Network inference: repeatability

The repeatability of CARNIVAL networks was evaluated by generating CARNIVAL-inferred networks for each dataset using four independent CPLEX random seeds inputs (i.e., random seed 1, 2, 3, and 4). The interactions and node attributes from each run were compared across all possible pairwise combinations to assess consistency across conditions. Interaction consistency was evaluated by checking that each interaction (i.e., node1 – interaction type – node2) was retained across runs. Edge weights were also compared, and discrepancies were quantified. If edge weights differed, the average and median differences were calculated for the pairwise comparison. Node consistency was assessed by comparing the presence of nodes across networks and evaluating differences in average node activity. If average activities were inconsistent, the difference was quantified, and the average and median differences were calculated across the pairwise comparison.

### Network inference: iterative network generation

To enhance the reproducibility of our network results, we applied an iterative approach to generate CARNIVAL networks. Each individual CARNIVAL network reached a near-optimal solution within 180 to 360 seconds (i.e., Gap = 0.00%). The optimal solution is assessed using the “Gap” output from the CPLEX optimizer, which represents the relative difference between the best feasible integer solution and the best known bound on the objective function. Despite the Gap reaching 0.00% within 360 seconds, CPLEX can continue running for many hours before confirming an optimal solution. To address possible network inference repeatability concerns and excessive computation times, we generated 30 iterations for each CARNIVAL network, using a CPLEX time limit of 600 seconds. This time limit ensured that the Gap reached 0.00%, indicating a near-optimal solution. While CPLEX operates deterministically, we introduced stochasticity by randomizing the order of input vectors (stdCARNIVAL: perturbation targets, TF activities, pathway scores; invCARNIVAL: TF activities, pathway scores) using the “sample” function in R. This approach ensured a more comprehensive evaluation of the full solution pool. The network solution pools from all 30 iterations were outputted and aggregated by averaging interaction weights and node attribute values to construct a final network. Some individual iterations showed a lack of network pruning, containing thousands of interactions instead of the expected 30 to 120. These outliers were removed before final aggregation to improve the quality of the resulting network.

### Network visualization

The CARNIVAL networks were visualized in Cytoscape (Shannon *et al*., 2003) by importing both the interaction file (weightedSIF) and the node attribute file (nodeAttributes). Network layouts were generated using the “yFiles Layout Algorithms”, specifically the “yFiles Hierarchic Layout”, following by the “yFiles Organic Edge Router”. In the final network visualizations, the node shape corresponds to nodeType: hexagons represent perturbation targets, diamonds represent inputted TFs, and circles represent inferred nodes. The node color reflects the node average activity, ranging from -100 to 100, where negative values (blue) indicate negative activity and positive values (red) indicate positive activity. Edge shading represents edge weight, ranging from 0 (white) to 100 (black). Target arrow shapes indicate regulatory effects: arrowheads denote stimulation, while flat-headed arrows denote inhibition.

### Numerical methods and software

R (version 4.3.2) was used to perform all network inference analyses. The IBM CPLEX optimization program (version 22.1.1) was used to solve the CARNIVAL integer linear programming problem. Cytoscape version 3.10.3 was used to perform all network visualization. The code generated for this study is freely available in the GitHub repository: https://doi.org/10.5281/zenodo.16907486.

## Results

### Transcriptional regulation of *RPS6KB1*

Using the OmniPath prior knowledge network (Ceccarelli *et al*., 2020), we identified 19 TFs that control *RPS6KB1* transcriptional activity (Figure 1). We then examined the regulatory roles of each TF in GeneCards (Stelzer *et al*., 2016), which revealed that 12 of the TFs regulate *RPS6KB1*. Among them, five primarily exert promoter-dominant regulation (*E2F1*, *ELF1*, *FOXA1*, *NRF1*, *UBTF*), while seven exhibit mixed promoter and enhancer regulation (*BATF*, *CEBPA*, *CTCF*, *EP300*, *GATA3*, *MYC*, *YBX1*; Figure 1) (Stelzer *et al*., 2016).

**Figure 1.**
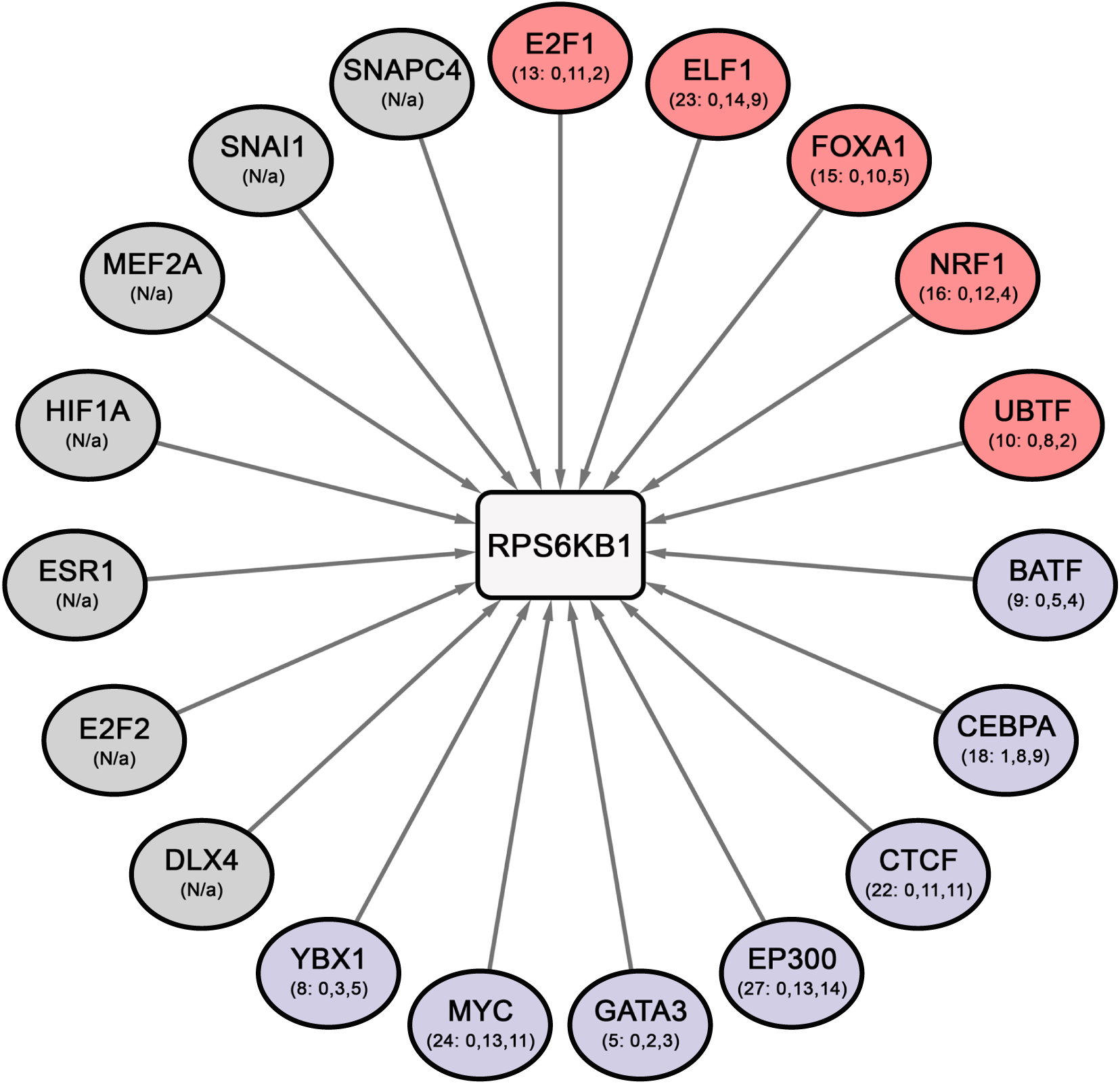
OmniPath-informed transcription factors (TFs) that control RPS6KB1 gene expression. The network was constructed by selecting the immediate upstream TFs of RPS6KB1 from the OmniPath signaling network. Each node displays TF binding site regulatory information extracted from GeneCards, with the four numbers in parentheses representing the (1) total number of TF binding sites, subdivided into those located in (2) the promoter region only, (3) both promoter and enhancer regions, and (4) the enhancer region only. “N/a” indicates that the TF was not documented in GeneCards for RPS6KB1. Node colors denote the predominant regulatory role of TF binding: red for TFs that bind primarily to promoter or promoter/enhancer regions, purple for TFs that bind approximately equally to promoter and enhancer regions, blue for TFs that bind primarily to enhancer regions, and gray for TFs absent from GeneCards. Notably, no TFs in GeneCards were reported to bind predominately to enhancer regions on RPS6KB1.

### Exercise dataset and differential gene expression analysis

In this study we used the data from Dickinson et al., which measured transcriptomic data from six recreationally active young men following acute aerobic and resistance exercise bouts. Transcript counts were measured at baseline, 1-hour, and 4-hours post-exercise (Supplementary Figure 1A). The raw transcript count data exhibited a bimodal distribution, with clear separation between the two modes at 0.5 log2(count). To reduce noise and improve the sensitivity of detecting differentially expressed genes, we excluded low counts [below 0.5 log2(count)] from further analysis (Sha *et al*., 2015). Variance stabilizing normalization was subsequently applied to the remaining transcript data (Supplementary Figure 1B).

Differential gene expression analysis was conducted to compare the baseline data with the 1-hour and 4-hour post-exercise data (Supplementary Figure 2). Minimal differential gene expression was observed between the baseline and 1-hour time point following both aerobic (Supplementary Figure 2A) and resistance exercise (Supplementary Figure 2C). In contrast, the 4-hour post-aerobic (Supplementary Figure 2B) and resistance exercise data (Supplementary Figure 2D) showed distinct differential gene expression compared to baseline. As a result, subsequent analyses focused on the baseline versus 4-hour post-resistance exercise datasets.

Next, we visualized the *RPS6KB1* transcript data to assess differential expression 4 hours following aerobic or resistance exercise (Figure 2). We observed increased *RPS6KB1* expression following resistance exercise, while no clear differential response was seen after aerobic exercise. However, the aerobic exercise data revealed participant-specific responses: participants 2 and 3 showed increased *RPS6KB1* expression, while participants 1 and 7 exhibited reduced expression.

**Figure 2:**
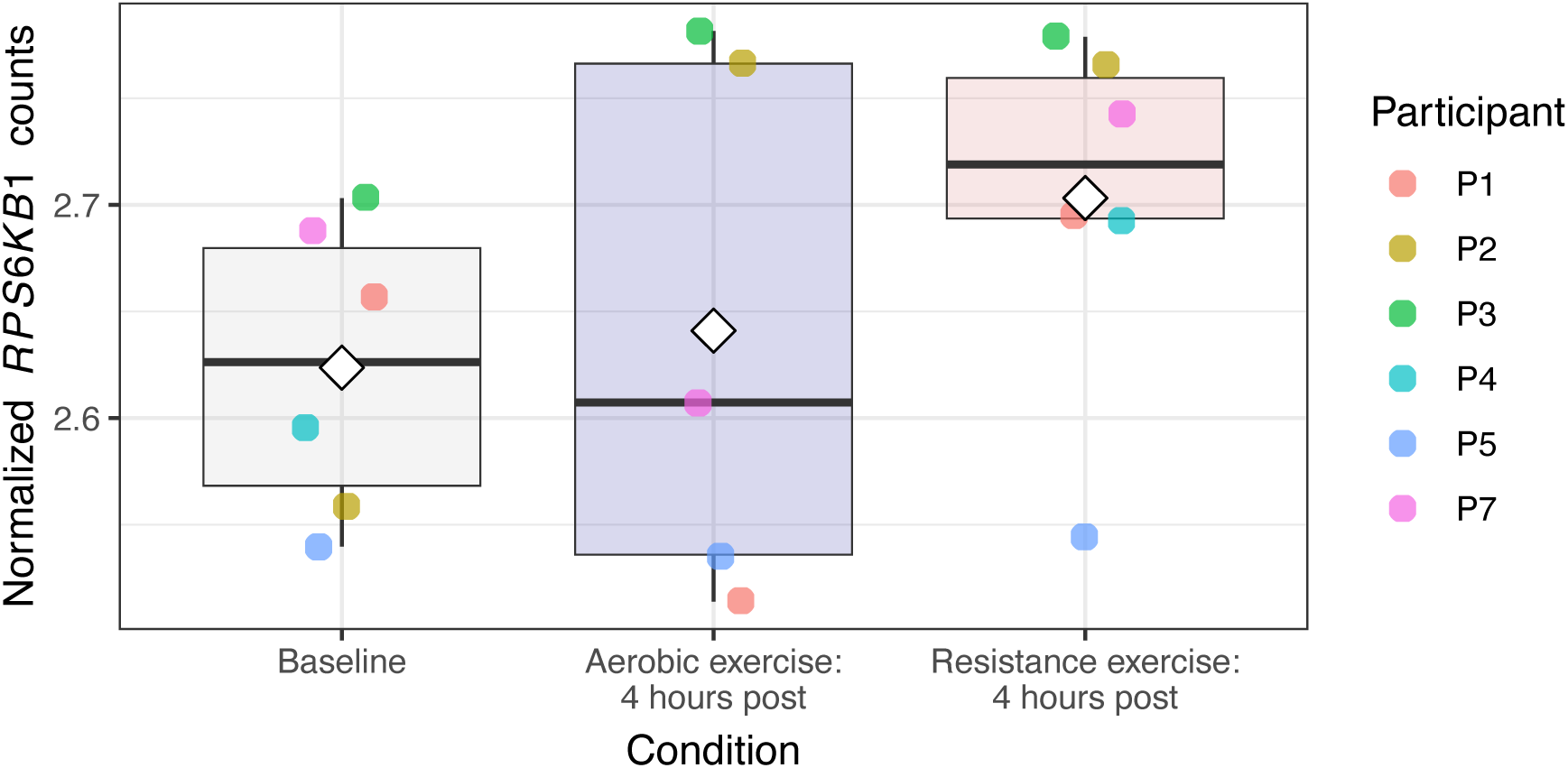
Normalized *RPS6KB1* transcript counts at baseline, 4 hours post-aerobic exercise, and 4 hours post-resistance exercise. Boxplots show the distribution of transcript counts for each condition, with the median represented by the thick black line, the first and third quartiles by the lower and upper hinges, and whiskers extending to the smallest and largest values within 1.5× the interquartile range. The mean is indicated by a diamond. Individual participant data points are overlaid to represent participant-specific responses.

### Transcription factor enrichment analysis

We estimated changes in the activities of TFs targeting *RPS6KB1* between baseline and 4 hours post-aerobic and resistance exercise using DoRothEA (Garcia-Alonso *et al*., 2019) and VIPER. Regulons with DoRothEA confidence scores of A, B, C, D, and E were included to estimate the NES and retain all TFs targeting *RPS6KB1*. Of the 19 TFs in Figure 1, activities were inferred for 17 (Figure 3) because neither *EP300* nor *UBTF* are classified as TFs in the DoRothEA database. *EP300* might be omitted from the database because it is a transcriptional co-activator (Gronkowska & Robaszkiewicz, 2024). The reason for the omission of *UBTF* is less clear.

**Figure 3:**
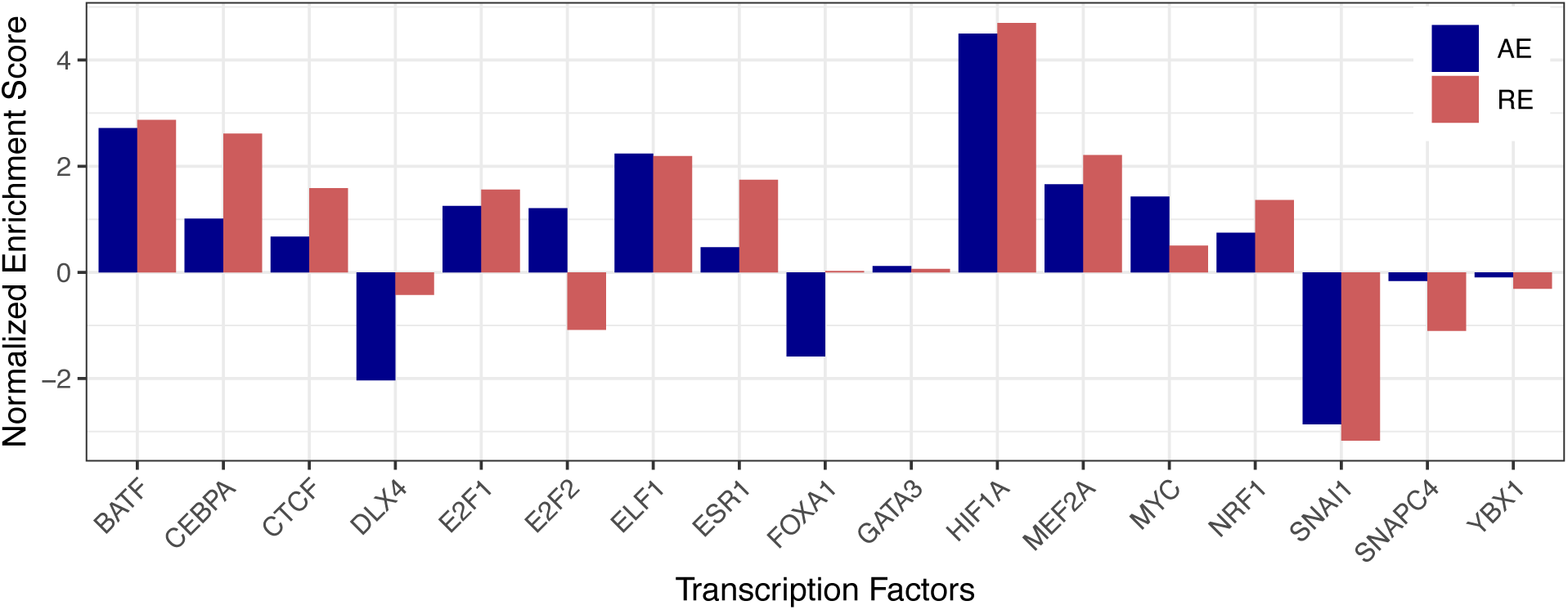
*RPS6KB1*-specific transcription factor activities. DoRothEA inferred normalized enrichment scores (NES) for the transcription factors regulating *RPS6KB1* gene expression 4 hours following aerobic or resistance exercise. The NES values were calculated using DoRothEA by comparing the baseline and 4-hours post-exercise time points.

We identified four TFs with high differential NES activity (i.e., NES > ±2) four hours post-aerobic and resistance exercise: *BATF* (AE: NES = 2.72, RE: NES = 2.87), *ELF1* (AE: NES = 2.24, RE: NES = 2.19), *HIF1A* (AE: NES = 4.50, RE: NES = 4.70), and *SNAI1* (AE: NES = -2.86, RE: NES = -3.17). In addition, we observed that most TFs responded similarly in magnitude and direction to both aerobic and resistance exercise. Notable exceptions were as follows: *DLX4* was strongly downregulated following only aerobic exercise (AE: NES = -2.03, RE: NES = -0.42), whereas *CEBPA* (AE: NES = 1.01, RE: NES = 2.62) and *MEF2A* (AE: NES = 1.66, RE: NES = 2.21) were strongly upregulated following resistance exercise. *E2F2* was moderately upregulated following aerobic exercise (NES = 1.21) and moderately downregulated following resistance exercise (NES = -1.08).

The TF activities were inferred based on the changes in expression levels of their target genes. To visualize the genes they regulate, we generated volcano plots of the highly differentially active TFs following both exercise conditions (Supplementary Figures 3 and 4).

### Pathway analysis

Pathway activities following both exercise bouts were inferred using PROGENy, which calculates pathway scores for 14 canonical pathways based on downstream transcriptional changes identified from perturbation experiments (Schubert *et al*., 2018). Pathway NES values between baseline and 4 hours post-aerobic and resistance exercise are presented in Figure 4. Both exercise conditions resulted in similar pathway responses, though resistance training generally induced a greater response. The NFκB (AE: NES = 7.08, RE: NES = 9.03) and TNFα (AE: NES = 6.20, RE: NES = 8.27) pathways were the most responsive following both aerobic and resistance exercise. To identify the main contributors to pathway enrichment, we generated scatterplots of the NFκB- and TNFα-responsive genes following both aerobic and resistance exercise (Supplementary Figures 5 and 6).

**Figure 4:**
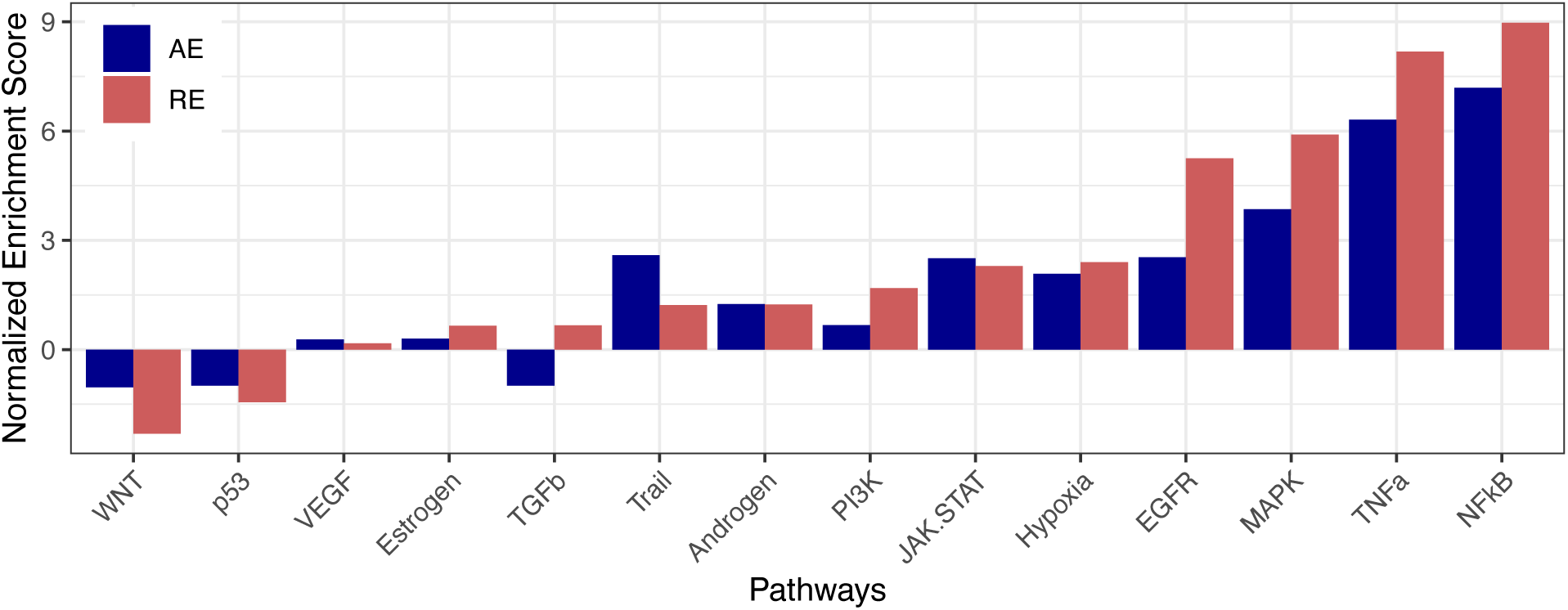
Pathway activities following aerobic and resistance exercise. Pathway activity normalized enrichment scores (NES) at 4-hours post-aerobic and resistance exercise. Relative pathway activity scores were calculated using PROGENy based on the differential gene expression between baseline and 4-hours post-exercise.

### Inferred signaling network

Using the inferred activities of the *RPS6KB1*-specific transcription factors and the predicted pathway activities, we applied standard CARNIVAL to infer the signaling network downstream of canonical exercise sensors following aerobic and resistance exercise.

We first evaluated the repeatability of the inferred standard CARNIVAL networks. Specifically, we performed four iterations of standard CARNIVAL for each of the aerobic and resistance exercise datasets, using the 50 canonical exercise sensors as targets of perturbation, the *RPS6KB1*-specific TFs, and a beta weight of 0.2 (Supplementary Tables 2-4). All iterations returned 100 solution pools, the maximum output allowed.

Across iterations, the inferred networks were consistent, with only mild differences in edge weights and average node activities that did not affect the biological interpretation of the results. Overall, the network topology, directionality of interactions, and relative node activities and edge weights were well preserved across runs, indicating strong repeatability of the inferred CARNIVAL solutions.

However, some minor differences were observed between individual runs. In the aerobic exercise data, three of the inferred networks lacked the *MAP3K7* node and the *MAP3K7*→*NFKB1* interaction, while one network pool lacked the *EGLN3* node and the *EGLN3*⊣*HIF1A* interaction. These interactions were included in the other networks, but with low edge weights. Differences in other values exhibited a balanced distribution between positive (activation) and negative (inhibition) discrepancies (Supplementary Tables 3-4).

Similarly, the resistance exercise networks revealed two discrepant interactions. Three of the networks included the *MAP3K8* node and the *MAP3K8*→*MAPK1* interaction, which were absent in one network. In contrast, the latter network included the *MAP3K1*→*MAPK1* interaction, which was missing from the previous three networks. The missing interactions had moderate edge weights in the networks where they were present.

Despite these minor differences, the resistance exercise networks preserved a similar percentage of interactions and nodes as the aerobic exercise networks (Supplementary Tables 3-4). However, there were more discrepancies in edge weights and node activities in the resistance exercise networks (aerobic exercise: 56 to 63% preserved edge weights, 62 to 69% preserved node activities; resistance exercise: 21 to 44% preserved edge weights, 41 to 59% preserved node activities; Supplementary Tables 3-4). Notably, the average and median absolute discrepancies in edge weights and node activities were smaller in the resistance exercise networks than the aerobic exercise networks. Again, the resistance exercise networks showed a relatively balanced distribution of positive (i.e., activation) and negative (i.e., inhibition) discrepancies (Supplementary Tables 3-4).

Overall, these differences were modest and did not meaningfully affect biological interpretation of the results. Nevertheless, given the inherent variability of single CARNIVAL network runs, we opted for an iterative approach to further improve repeatability. Specifically, we generated 30 iterations for each CARNIVAL network and aggregated the results to construct the final networks.

### Aerobic and resistance exercise-mediated control of *RPS6KB1*

The stdCARNIVAL-inferred networks for both aerobic (Figure 5) and resistance exercise (Figure 6) identified ten *RPS6KB1*-specific TFs and eleven canonical exercise sensors that featured a mix of intracellular sensors and hormone receptors. While the networks for both exercise modalities shared many of the TFs (*CTCF*, *ELF1*, *E2F1*, *E2F2*, *HIF1A*, *MEF2A*, *NRF1*, *SNAI1*), only the aerobic exercise network included *FOXA1* and *MYC*, whereas only the resistance exercise network contained *CEBPA* and *ESR1*. Differences also emerged in the inferred exercise sensors. The aerobic exercise network contained eight intracellular sensors (*EGLN1*, *EGLN2*, *EGLN3*, *HIF1AN*, *MAP3K7*, *PRKAA1*, *PRKACA*, *SIRT1*) and three hormone receptors (*ACVR2B*, *INSR*, *TGFBR1)*, while the resistance exercise network featured seven intracellular sensors (*MAP3K1*, *MAP3K2*, *MAP3K7*, *MAP3K8*, *PRKAA1*, *PRKACA*, *SIRT1*) and four hormone receptors (*ACVR2B*, *AR*, *INSR*, *TGFBR1*). The downstream TF controlled by each exercise sensor (Supplementary Table 5) and exercise sensors that control each TF (Supplementary Table 6) were analyzed to highlight key regulatory mechanisms.

**Figure 5:**
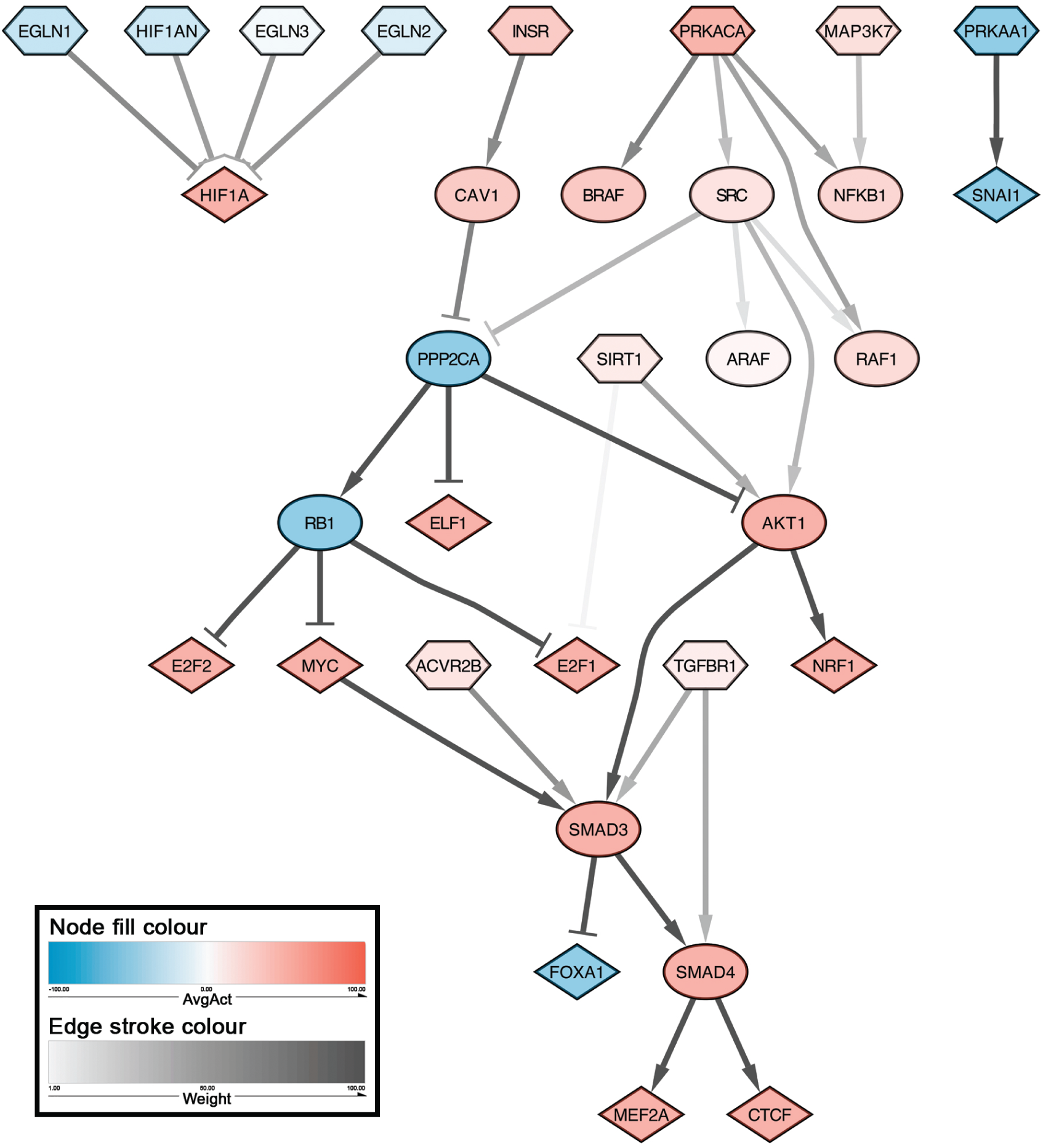
Standard CARNIVAL-inferred network of *RPS6KB1*-specific TF control by exercise sensors following aerobic exercise. Network visualization details are provided in the Methods.

**Figure 6:**
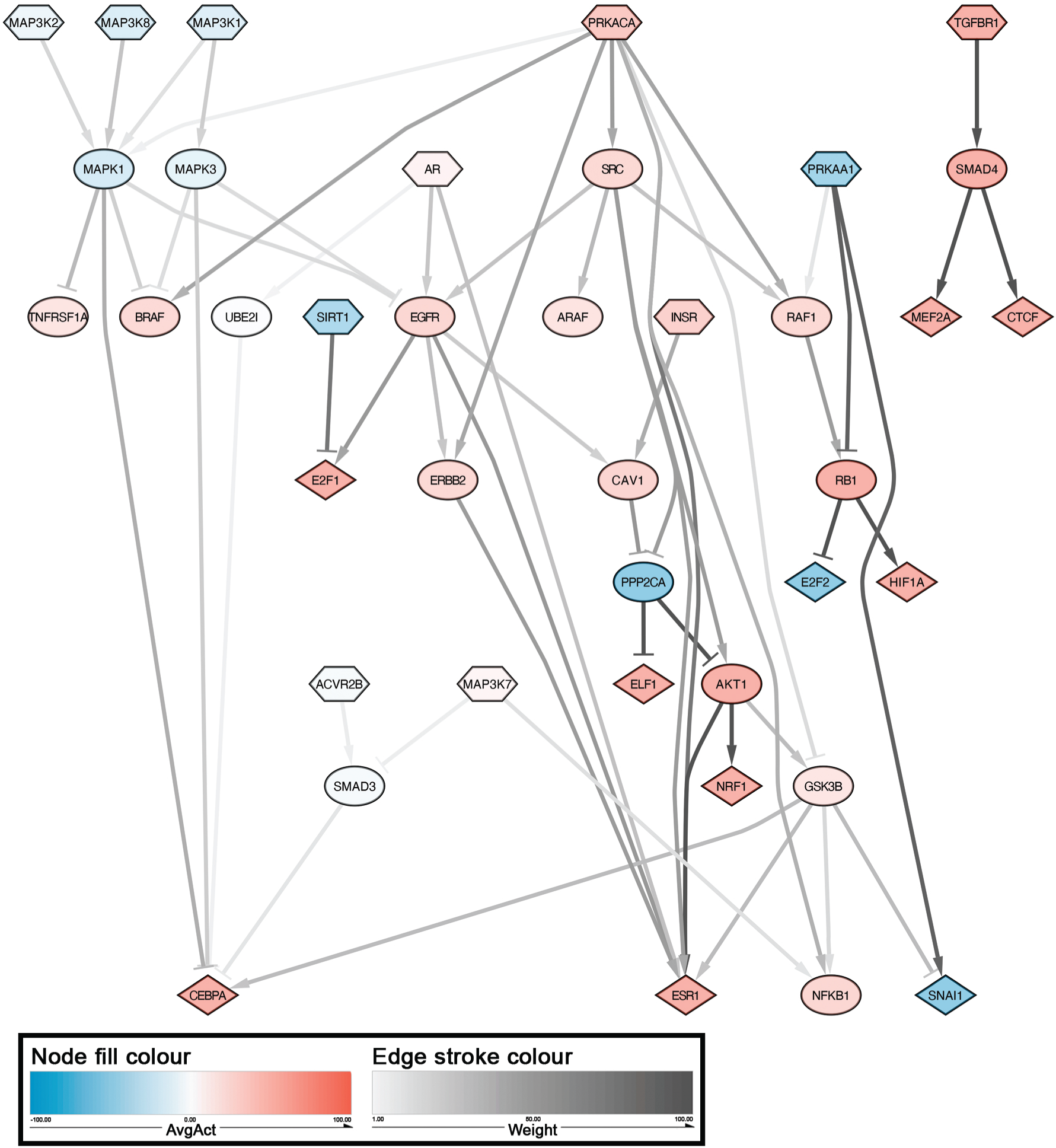
Standard CARNIVAL-inferred network of *RPS6KB1*-specific TF control by exercise sensors following resistance exercise. Network visualization details are provided in the Methods.

Among the TFs with high differential activity (Figure 3), three from aerobic exercise – *ElF1*, *HIF1A*, and *SNAI1* – were incorporated into the inferred network, along with five from resistance exercise – *CEBPA*, *ELF1*, *HIF1A*, *MEF2A*, and *SNAI1* (Figures 5 and 6). In the aerobic-exercise-responsive network, *ELF1* was regulated by *INSR* via *CAV1*/*PPP2CA* and *PRKACA* through *SRC*/*PPP2CA*, while *HIF1A* was jointly regulated directly by *EGLN1*, *EGLN2*, *EGLN3*, and *HIF1AN*. *SNAI1* was consistently regulated directly by *PRKAA1* across the solution pool. In the resistance-exercise-responsive network, *CEBPA* was jointly regulated by *MAP3K1*, *MAP3K2*, and *MAP3K8* via *MAPK1* and *MAPK3. ELF1* was predominantly regulated by *PRKACA* through *SRC* or the *SRC*/*EGFR*/*CAV1*/*PPP2CA* axis, with additional regulation by *INSR* via *CAV1*/*PPP2CA* and moderate input from *AR* through the *EGFR*/*CAV1*/*PPP2CA* pathway. *HIF1A* was primarily regulated by *PRKAA1* via *RB1*, with additional regulation by *PRKACA* through *RAF1*. *MEF2A* was consistently regulated across the solution pool by *TGFBR1* via *SMAD4*. *SNAI1* was predominantly regulated directly by *PRKAA1*, with moderate contributions from *INSR* via the *CAV1*/*PPP2CA*/*AKT1*/*GSK3B* axis.

To identify alternative regulatory mechanisms, we applied invCARNIVAL to infer additional exercise sensors influencing TFs targeting *RPS6KB1* (Figure 7). The aerobic-exercise-responsive network, based on 29 iterations (excluding one outlier), included 10 *RPS6KB1*-specific TFs and 17 sensor nodes immediately downstream of the perturbation. Seven sensor nodes had very low average activity (below 1) and were removed from the final network. Several remaining low-activity sensors converged on the protein phosphatase 2A subunits, *PPP2A* and *PPP2B*, indicating a possible redundancy in TF regulation. Notably, *TFDP2* and *TLX1* displayed higher activity, indicating stronger inferred regulatory roles. *TFDP2* directly regulated *E2F2*, while *TLX1* influenced several TFs, including *CTCF*, *E2F1*, *FOXA1*, *MYC*, and *SNAI1*, through *RB1*.

**Figure 7:**
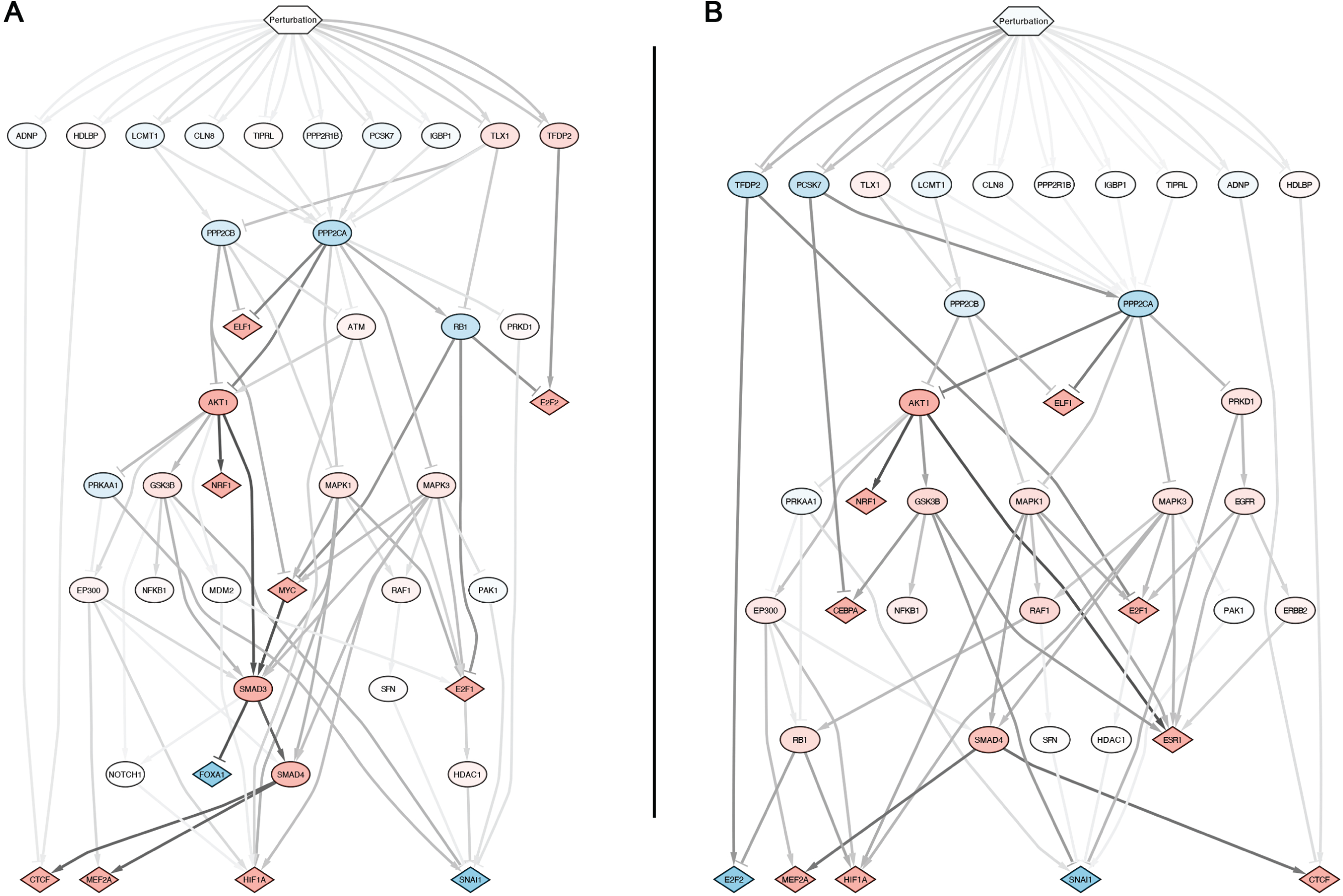
Inverse CARNIVAL-inferred networks. The networks represent the regulation of *RPS6KB1*-specific TFs following (A) aerobic and (B) resistance exercise. Network visualization details are provided in the Methods.

The resistance-exercise-responsive network derived from 28 iterations (excluding two outliers), included 10 *RPS6KB1*-specific TFs and 10 sensor nodes immediately downstream of the perturbation. Similar to the aerobic exercise network, many sensors exhibited low activity, suggesting weak or redundant regulatory influence. Several nodes converged on *PPP2A*, reinforcing its role as a central regulator. Comparatively, *PCSK7*, *TFDP2*, and *TLX1* showed higher inferred activities. These nodes are inferred to primarily act on the protein phosphatase 2A subunits (*PPP2CA* and *PPP2CB*), both of which share similar downstream effects and regulate all ten TFs included in the network. Further, *PCSK7* and *TFDP2* were inferred to have direct regulatory interactions with specific TFs: *PCSK7* targeted *CEBPA*, while *TFDP2* influenced *E2F1* and *E2F2*.

## Discussion

In this study, we applied the causal network inference approach CARNIVAL to map the exercise-driven signaling events controlling *RPS6KB1* transcriptional activity. While previous bioinformatics approaches in exercise biology – such as differential expression and enrichment analyses, pathway-level inference, network-based methods, and meta-analysis approaches – have provided valuable insights into the transcriptional control of genes, these methods are largely descriptive and offer limited ability to infer upstream regulatory mechanisms. By integrating prior knowledge of signaling networks with transcriptomic data through causal reasoning, CARNIVAL infers both the directionality and causality of upstream signaling events controlling observed transcriptional responses. Applied here, CARNIVAL provides a unique framework to identify potential exercise-specific interventional targets by mapping how upstream signals propagate to downstream transcriptional events.

Our analyses revealed two key findings. First, AE and RE activated most TFs controlling *RPS6KB1* in a similar magnitude and direction, with the exceptions of *DLX4*, which responded only to AE, and *CEBPA* and *MEF2A*, which only responded to RE. Second, the inferred networks indicate that AMPK, protein kinase A, and insulin receptor may be central exercise sensors controlling many TFs following both AE and RE. However, AE uniquely engaged hypoxia-related sensors in regulating *HIF1A*, while RE distinctly controlled *MEF2A* through TGF-β signaling, *CEBPA* via MAP3Ks, and *E2F1* via sirtuin. Together, these results motivate new hypotheses regarding the molecular control of *RPS6KB1* transcription following aerobic and resistance exercise.

### Main findings and their implications

*ESR1* is one of the few TFs that has been researched for its role in controlling *RPS6KB1* expression by binding to its promoter region (Maruani *et al*., 2012; Berman *et al*., 2017). *ESR1* encodes the ERα protein, an estrogen-dependent TF that controls many genes across many tissues (Hevener *et al*., 2020). In skeletal muscle, ERα controls muscle force and endurance by modulating mitochondrial function (Hevener *et al*., 2020; Yoh *et al*., 2022, 2023), metabolic homeostasis and insulin action (Hevener *et al*., 2020), and controlling satellite function and muscle regeneration (Collins *et al*., 2019). ERα is highly expressed in skeletal muscle and its levels inversely correlate with metabolic health (i.e., lower ERα levels have been observed in healthy mice) (Ribas *et al*., 2016).

Our results show that ERα was modestly activated following AE and upregulated following RE, primarily through signaling downstream of *PRKACA*, the gene encoding protein kinase A (PKA). PKA is a sensor of cyclic AMP (cAMP), a secondary messenger produced intracellularly downstream of adenyl-cyclase-mediated conversion of ATP in response to hormones and neurotransmitters (Berdeaux & Stewart, 2012; Sassone-Corsi, 2012); cAMP accumulates following maximal cycling exercise (Chasiotis *et al*., 1983). PKA phosphorylates several residues on ERα, including S236, S305, S338, and S518 (Chen *et al*., 1999; Lazennec *et al*., 2001). Functionally, the phosphorylation of S236 in fibroblast-like cells inhibits ERα dimerization and reduces its DNA binding in the absence of estrogen (Chen *et al*., 1999). However, in the presence of estrogen, PKA-induced phosphorylation of ERα increases its transactivational function (Chen *et al*., 1999). Given the estrogen-dependent activation of ERα by PKA signaling, the inferred activation of ERα in our results suggests a potential estrogen-dependent mechanism. Estradiol, a form of estrogen, increases following resistance exercise in both males and females, but at reduced magnitudes and lower absolute concentrations in men (Copeland *et al*., 2002; Wolf *et al*., 2012). The exercise-mediated increase in estradiol in males suggests a viable mechanism for the inferred PKA-mediated expression of ERα, with the data used for network generation exclusively featuring male participants.

*SNAI1*, a *RPS6KB1*-specific TF, was highly downregulated following both AE and RE. *SNAI1* encodes the Snail1 protein, a member of the Snail family of zinc-finger transcription factors, that plays a key role in epithelial-to-mesenchymal cell transition (EMT) (Kaufhold & Bonavida, 2014). EMT is a biological process essential for embryo formation, wound healing, and tissue regeneration, during which epithelial cells lose their apical polarity and adopt mesenchymal characteristics (Kaufhold & Bonavida, 2014; Kielbik *et al*., 2024). Snail1 facilitates EMT by repressing E-cadherin and activating signaling pathways (Kaufhold & Bonavida, 2014; Kielbik *et al*., 2024). While Snail1 is expressed in human skeletal muscle, its levels are lower compared to heart, placenta, and lung (Paznekas *et al*., 1999). Notably, Snail1 expression increases following drug-induced muscle injury during muscle regeneration (Elia *et al*., 2022).

Previous experimental findings suggest a bidirectional regulatory relationship between p70S6K and Snail1. In ovarian cancer cells, constitutively active p70S6K induced transcriptional upregulation of Snail, which repressed levels of E-cadherin (Pon *et al*., 2008). Reciprocally, Snail1 controlled p70S6K levels through a miR-128-dependent mechanism in prostate cancer cells (Tao *et al*., 2014). Depletion of Snail1 increased miR-128 expression, which subsequently reduced p70S6K levels, inhibited cell growth, and reduced glucose consumption and lactate production (Tao *et al*., 2014). Our results indicate that Snail1 expression is regulated by *PRKAA1*, which encodes AMPK α-subunit 1, following both AE and RE. Indeed, prior research demonstrates that AMPK interacts with Snail1, but contradictory directions of influence have been observed. In cancer cells, activation of the AMPK heterotrimer increases Snail1 expression, whereas inhibition of AMPK represses Snail1 expression (Saxena *et al*., 2018) – this positive regulation is consistent with the inferred networks. Conversely, following glucose deprivation in breast cancer cells, AMPKα phosphorylated Snail1 at Ser11, which targeted Snail1 for degradation (Li *et al*., 2024). This AMPKα-mediated negative control of Snail1 contradicts our inferred network. Despite the uncertainty by which AMPK controls Snail1 expression, clear evidence exists for its regulation of the Snail1 and for Snail1-mediated control of p70S6K levels, albeit through miR-128 mechanisms.

*HIF1A*, a *RPS6KB1*-specific TF, was inferred to be the most differentially activated TF following both AE and RE (Figure 3). *HIF1A* encodes the alpha subunit of hypoxia-inducible factor 1 (HIF-1) (Iyer *et al*., 1998). HIF-1 is a heterodimer transcription factor consisting of HIF-1α and HIF-1β/ARNT (aryl hydrocarbon receptor nuclear translocator) subunits, and is considered a critical regulator of cellular oxygen detection and adaptation (Choudhry & Harris, 2018). HIF-1α is highly expressed in skeletal muscle in normoxic and hypoxic conditions (Stroka *et al*., 2001), and controls the expression of genes involved in oxygen consumption, erythropoiesis, angiogenesis, mitochondrial metabolism, and glucose metabolism (Favier *et al*., 2015; Choudhry & Harris, 2018). In normoxia, HIF-1α is hydroxylated by prolyl hydroxylase domain (PHD) enzymes and factor inhibiting HIF (FIH), leading to its proteasomal degradation or transcriptional inhibition, respectively (Favier *et al*., 2015). Under hypoxic conditions, PHDs become inactivated, leading to HIF-1α stabilization and its dimerization with HIF-1β (Choudhry & Harris, 2018). The HIF-1 heterodimer translocates to the nucleus, where it binds to hypoxia response elements in the promoter region of hypoxia-responsive genes (Choudhry & Harris, 2018). Muscle hypoxia can occur during both aerobic (Boone *et al*., 2016) and resistance exercise (Pereira *et al*., 2007; de Oliveira *et al*., 2020) because both exercise types reduce intracellular PO_2_, thus promoting HIF-1 activity.

Research in brain tumors has shown that *RPS6KB1* expression is associated with hypoxia-induced genes, implying a potential link between *RPS6KB1* and *HIF1A* (Ismail, 2012). Although a direct interaction between *HIF1A* and *RPS6KB1* has yet to be established, evidence supports a functional relationship between hypoxia and increased *RPS6KB1* expression. Interestingly, this relationship appears to be bidirectional, as HIF-1α levels have been reported to depend on the phosphorylation status of p70S6K (Liu *et al*., 2009). Our results indicate that *HIF1A* activity is differentially controlled following acute AE and RE. Following AE, *HIF1A* was jointly controlled by *EGLN1*, *EGLN2*, *EGLN3*, and *HIF1AN*, which encode PHD1-3 and FIH, respectively. These proteins are well-established negative controllers of HIF-1α under normoxic conditions (Favier *et al*., 2015; Choudhry & Harris, 2018). In our inferred regulatory network, PHD1-3 and FIH showed reduced inhibitory activity towards HIF-1α, consistent with the hypoxic intramuscular environment induced by AE (Boone *et al*., 2016). In contrast, following RE, *HIF1A* was jointly controlled through signaling downstream of *PRKAA1* (AMPK α-subunit 1) and *PRKACA* (PKA). Indeed, cultured glioblastoma cells deprived of serine and glycine exhibited increased AMPK kinase activity and AMPK-mediated HIF-1α stabilization and transactivation (Yun *et al*., 2023). Similarly, PKA activity can enhance HIF-1α transcriptional activity directly by phosphorylating its Thr63 and Ser692 residues (Bullen *et al*., 2016) and indirectly via Raf-1 signaling (Häfner *et al*., 1994; Dhillon *et al*., 2002; Dumaz & Marais, 2003; Lim *et al*., 2004). However, literature suggests that PKA inhibits Raf-1, which contradicts our inferred network (Häfner *et al*., 1994; Dumaz & Marais, 2003). Nevertheless, the RE network suggests that HIF-1α is primarily controlled via AMPK signalling. Together, these networks highlight exercise-specific control of HIF-1α and predict that hypoxia-induced signaling functionally influences *RPS6KB1* expression.

### Limitations

Four noteworthy limitations may impact the results of this study and their interpretation. First, limitations may exist with the dataset used in our analysis. The dataset featured a small sample of six untrained male participants who completed both AE and RE in a crossover design. While this design helps control for interindividual variability, the small sample size limits the generalizability of the findings. Future studies could compare our results with the MetaMEx database (Pillon *et al*., 2020) to assess the generalizability of the transcriptional responses or incorporate meta-analyzed transcriptomic data instead of data from single studies. Additionally, the transcriptomic response was measured following a single, acute exercise bout, which may induce a disproportionate inflammatory response (Cerqueira *et al*., 2020). This response could obscure the exercise-specific transcriptomic response typically observed after training (Thomas *et al*., 2024). For example, the signaling response tends to be elevated after an initial acute exercise bout but attenuates with subsequent sessions following adaptation (Langer *et al*., 2022). Thus, the transcriptomic response to future sessions may differ from the one featured in this dataset. Lastly, our inferred network reflects the 4-hours post-exercise state. Although robust transcriptional responses are typically detected at this time (Yang *et al*., 2005; Louis *et al*., 2007), exercise induces sequential “waves” of gene expression, encompassing both early and late transcription events (Raue *et al*., 2015; Pillon *et al*., 2022). Consequently, transcriptional events occurring outside the 4-hour window may not have been captured.

Second, the inferred networks used in this study – the TF-target network for *RPS6KB1* and the protein-protein interaction network for CARNIVAL – are inherently biased towards known biology (Liu *et al*., 2019). Consequently, novel molecular interactions or alternative interpretations of existing interactions could alter the network results and study interpretations. Additionally, potential errors in interaction annotations, including incorrect directional relationships (i.e., stimulatory or inhibitory), could similarly affect network inference and interpretation.

Third, the interpretation of the inferred networks is constrained by the available literature. To our knowledge, research on the transcriptional control of *RPS6KB1* is limited, and its control in skeletal muscle remains unexplored. Much of the existing evidence linking the identified TFs to *RPS6KB1* control is derived from studies of cancer cells. While the fundamental mechanisms for transcriptional control are often conserved across tissues, their function and expression levels may differ. For example, proteins can be expressed at markedly different levels in cancer cells, which may drive responses not normally seen in healthy tissue (Pessoa *et al*., 2022). Regardless, this study aimed to generate strong hypotheses for the exercise-mediated control of *RPS6KB1*, which can ultimately only be informed based on available evidence. Our findings featuring inferred causal networks motivate specific experimentally testable hypotheses regarding the control of *RPS6KB1* in skeletal muscle.

Lastly, the broader implications of the findings presented in this study deserve consideration. Here, we have inferred the TF-mediated control of *RPS6KB1* gene expression. While TF activity is essential for initiating the production of new functional proteins, transcriptional control represents only the first step in this multi-step process. TF-mediated gene expression leads to the generation of new mRNA transcripts, which may subsequently be translated into proteins such as p70S6K. However, post-transcriptional control and post-translational modifications further influence protein synthesis and function. Regarding post-transcriptional control, non-coding RNAs such as microRNA (miRNA) and long non-coding RNA (lncRNA) play key roles (Ratti *et al*., 2020; Nie *et al*., 2023). MicroRNA are short RNA molecules that inhibit translation by binding to target mRNAs, leading to transcript degradation or translational inhibition (Ratti *et al*., 2020; Nie *et al*., 2023). Long non-coding RNA, which are much longer RNA transcripts of more than 200 nucleotides in length, serve several functional roles (Ratti *et al*., 2020; Nie *et al*., 2023). Notably, lncRNAs can function as competitive endogenous RNAs, sequestering miRNAs and thereby relieving their inhibitory effects, ultimately enhancing translation (Ratti *et al*., 2020; Nie *et al*., 2023). Therefore, the post-transcriptional control mediated by the interplay between miRNA and lncRNA adds further complexity to the control of p70S6K levels beyond TF-mediated transcriptional control.

### Future directions

Several unresolved questions remain regarding the transcriptional control of *RPS6KB1* that warrant further investigation. Among the TFs discussed, *ESR1* is the most studied controller of *RPS6KB1*, making evaluation of its role in skeletal muscle a particularly promising avenue for modulating *RPS6KB1* expression. Because *ESR1* expression is mediated by PKA activity in an estrogen-dependent manner, it is worth exploring whether targeted *ESR1* activation elicits sex-specific muscle responses. Here, the network motivates the hypothesis that PKA activation in females may induce greater increases in *RPS6KB1* expression. Furthermore, the transcriptional factor Snail1 requires experimental study in skeletal muscle to clarify how it is controlled by AMPK following exercise. Given Snail1’s strong differential response to exercise, it will be important to determine whether it directly controls *RPS6KB1*. Similarly, the relationship between hypoxia and *RPS6KB1* remains poorly defined. Establishing whether this is mediated through *HIF1A* is of particular interest, because it would provide a direct mechanistic link between exercise-induced hypoxia and *RPS6KB1* expression. Finally, a critical next step is to assess the extent to which the predicted changes in *RPS6KB1*-specific TFs translate into measurable increases in *RPS6KB1* mRNA and p70S6K protein abundance.

## Conclusions

In summary, we identified plausible TFs that control *RPS6KB1* expression using a curated database and inferred networks linking these TFs to canonical intracellular exercise sensors and hormone receptors following AE and RE. Our findings reveal that while many *RPS6KB1*-specific TFs respond similarly to acute AE and RE, some exhibit distinct responses depending on exercise modality. Our inferred networks and preliminary interpretation highlight energy balance pathways, second messenger signaling, and hormonal regulation as key controllers of p70S6K levels. Should the identified exercise-responsive sensors and pathways controlling *RPS6KB1* in this study be experimentally verified, then the molecules could serve as targets for pharmaceutically manipulating p70S6K levels.

## Data availability statement

The R code generated in this study is available in the GitHub repository: https://doi.org/10.5281/zenodo.16907486.

## Author contributions

T.J.M.: Conceptualization, Data curation, Formal analysis, Investigation, Methodology, Software, Visualization, Writing – original draft, Writing – review and editing.

R.Z.: Data curation, Formal analysis, Investigation, Software, Visualization, Writing – review and editing.

A.D.: Conceptualization, Methodology, Software, Supervision, Writing – review and editing;

J.S.R.: Resources, Project administration, Supervision, Writing – review and editing;

D.C.C.: Conceptualization, Funding acquisition, Project administration, Resources, Supervision, Writing – review and editing.

## Acknowledgements

We thank Drs. Eldon Emberly and Daniel Moore for their detailed feedback on this work and Dr. Eldon Emberly for critically proofreading an earlier version of the manuscript.

