## Supplementary Information for "Bioinformatic inference of the exercise-responsive control of p70 S6 kinase through *RPS6KB1* expression"

### Bioinformatic inference of the exercise-responsive p70 S6 kinase signalling network

#### 1 Supplementary Figures and Tables

##### 1.1 Supplementary Figures

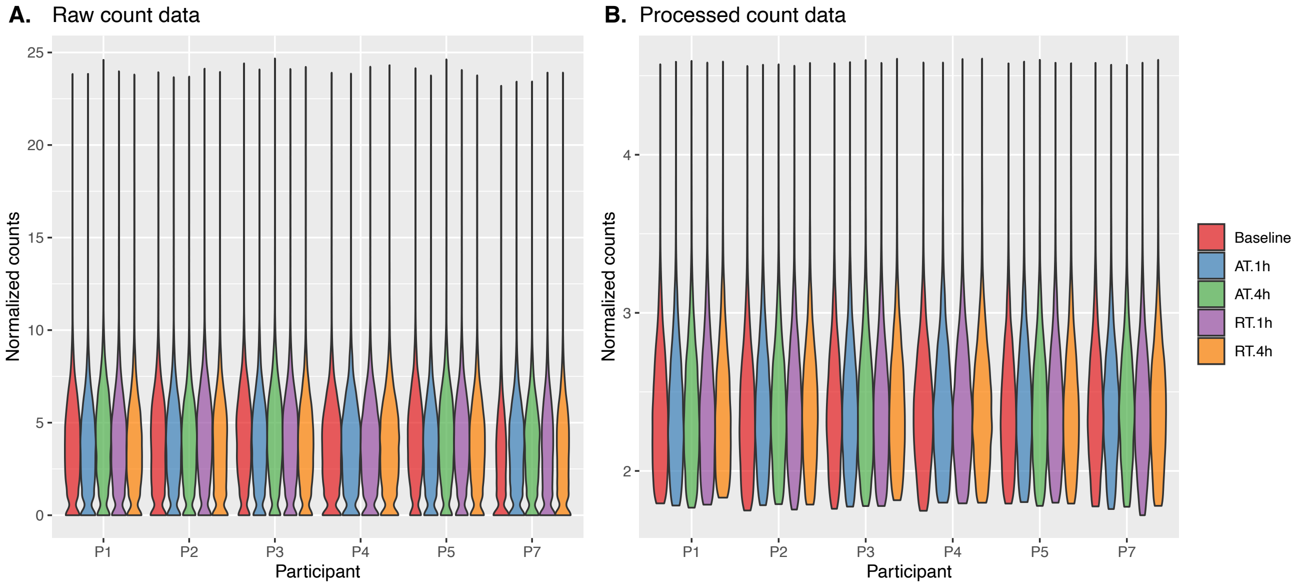


**Supplementary Figure 1.** Violin plots of (A) log2-transformed raw gene count data and (B) processed gene count data for the six participants (P1-5, P7) at baseline, 1-hour (1h) or 4-hour (4h) post-aerobic exercise, and 1-hour or 4-hour post-resistance exercise. The raw count data exhibited a bimodal distribution, with low-count genes below 0.5 log2(count) forming one mode. These low-count genes were removed during data processing. Variance stabilizing normalization was subsequently applied to generate the processed gene count data shown in panel B.

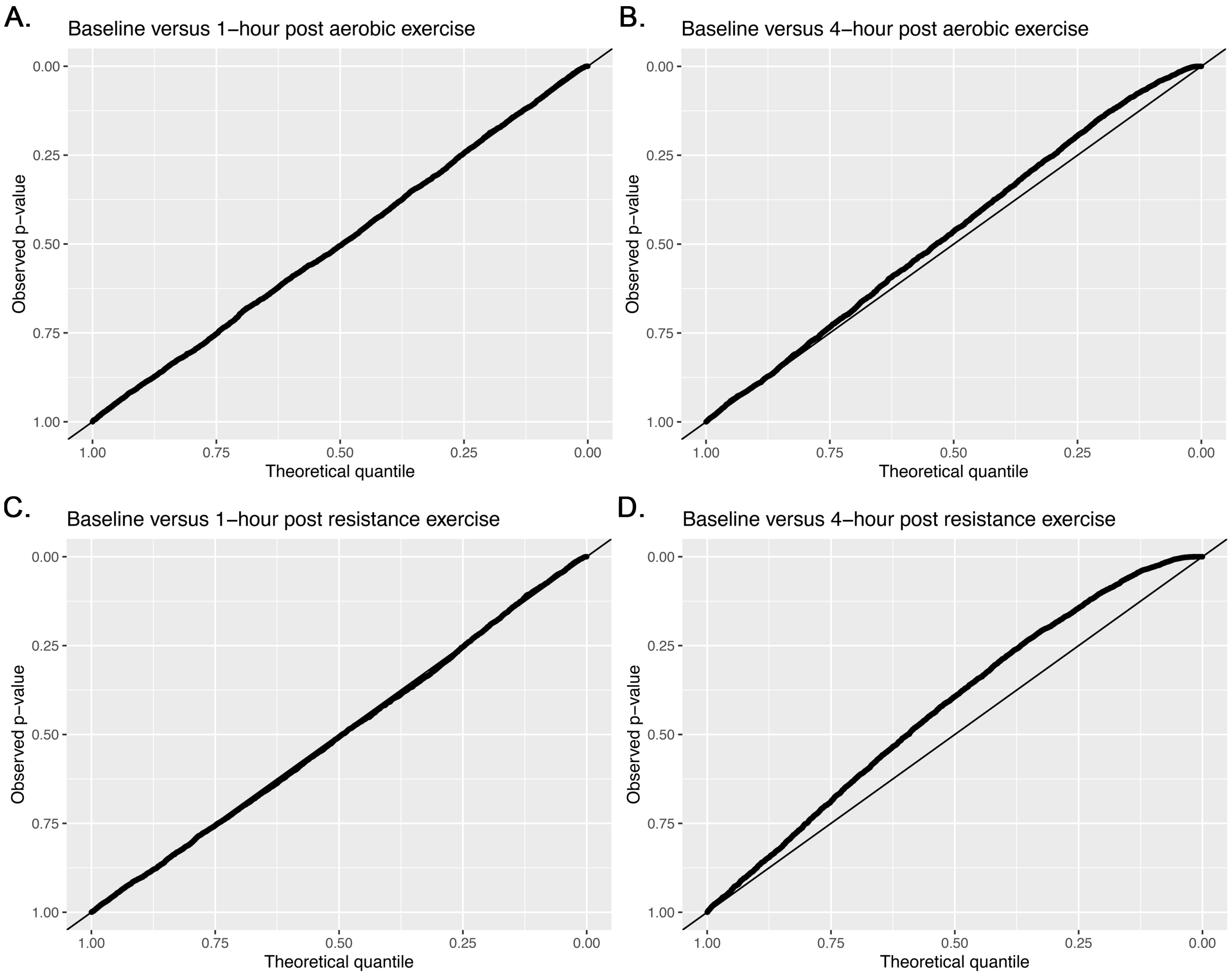


**Supplementary Figure 2.** Quantile-quantile plots between the observed p-value distribution from the differential gene expression analysis comparing the baseline condition with (A) the 1-hour post aerobic exercise condition, (B) the 4-hour post aerobic exercise condition, (C) the 1-hour post resistance exercise condition, and (D) the 4-hour post resistance exercise condition. The observed p-values are plotted against the expected p-values under the null distribution. The line of identity is shown, with deviations from this line indicating enrichment of differentially expressed genes. Greater deviations reflect stronger evidence for differential gene expression.

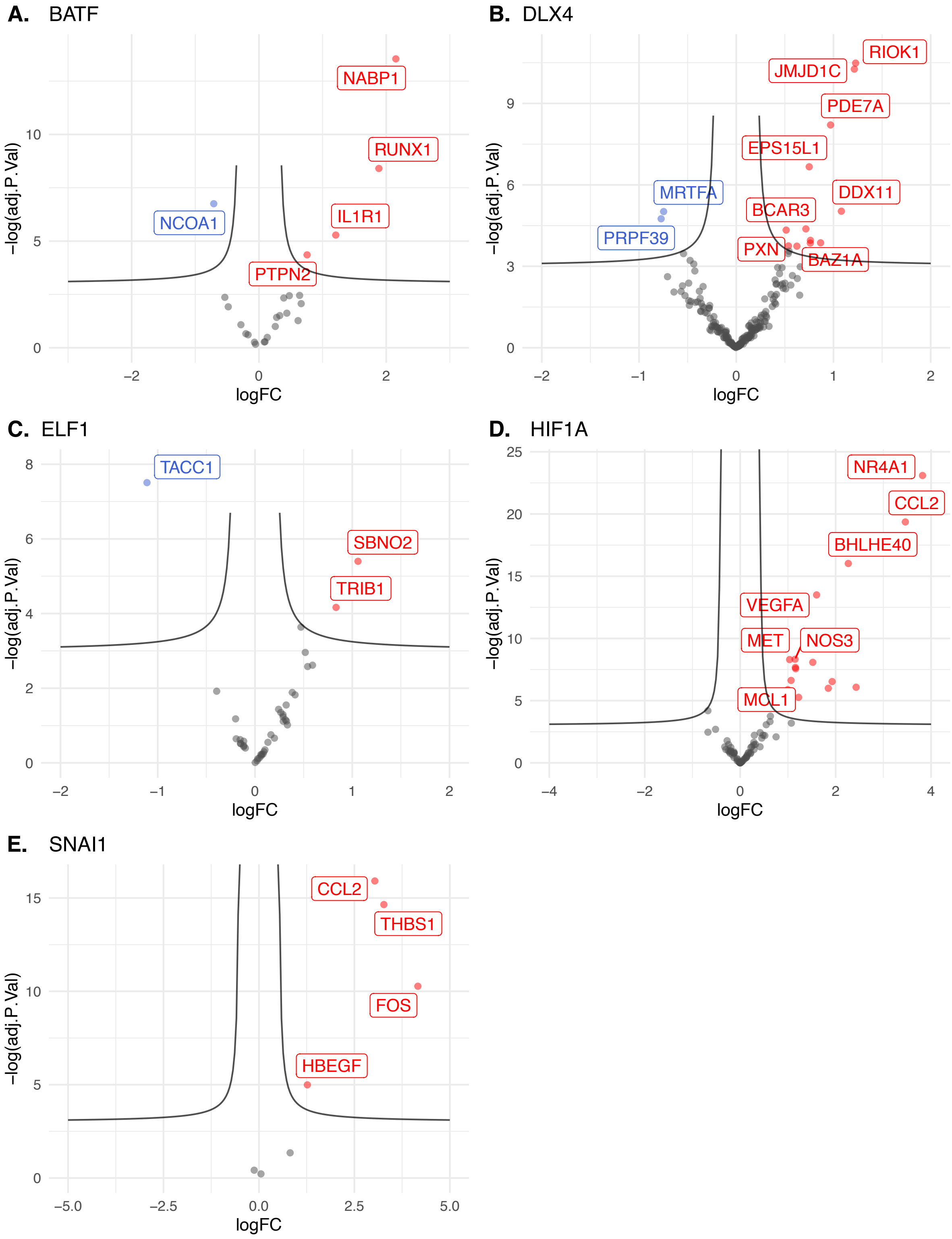


**Supplementary Figure 3.** Volcano plot of target gene expression levels for highly differentially active transcription factors 4 hours after the aerobic exercise bout: (A) *BATF*, (B) *DLX4*, (C) *ELF1*, (D) *HIF1A*, and (E) *SNAI1*. Genes with greater upregulation or downregulation are positioned toward the right or left, respectively, while genes with higher statistical significance are closer to the top. Blue points represent downregulated genes, red points represent upregulated genes. logFC = log fold change.

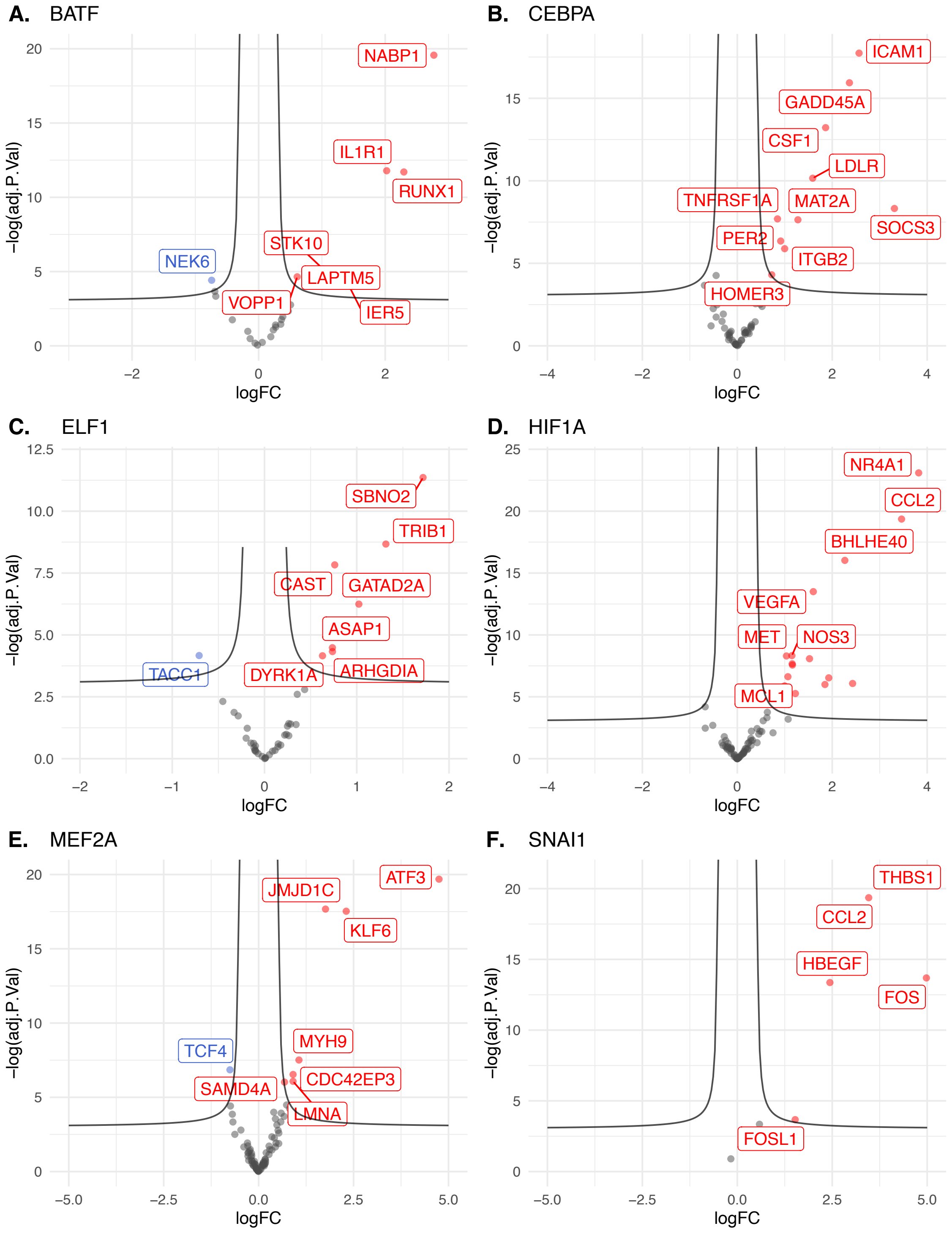


**Supplementary Figure 4.** Volcano plot of target gene expression levels for highly differentially active transcription factors 4 hours after the resistance exercise bout: (A) *BATF*, (B) *CEBPA*, (C) *ELF1*, (D) *HIF1A*, (E) *MEF2A*, and (F) *SNAI1*. Genes exhibiting greater upregulation or downregulation are positioned towards the right or left, respectively, while genes with higher statistical significance are closer to the top. Blue points represent down regulated genes, red points represent upregulated genes. logFC = log fold change.

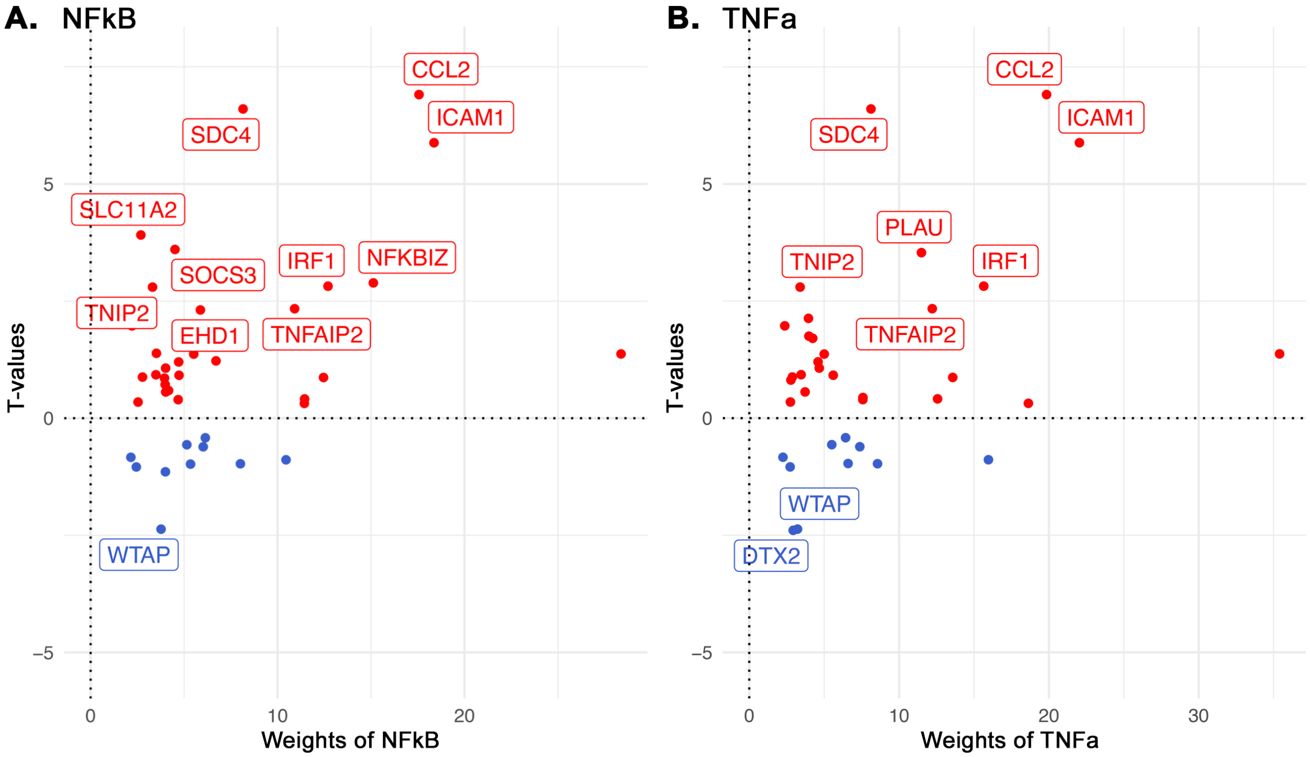


**Supplementary Figure 5.** Scatterplots of (A) NFkB and (B) TNFa pathway-responsive genes 4-hours following aerobic exercise. Pathway activity scores were calculated using PROGENy showing the association between pathway-responsive gene expression and the predicted activity of the signalling pathways. Red points indicate genes contributing positively to the pathway score, while blue points indicate genes contributing negatively to the pathway.

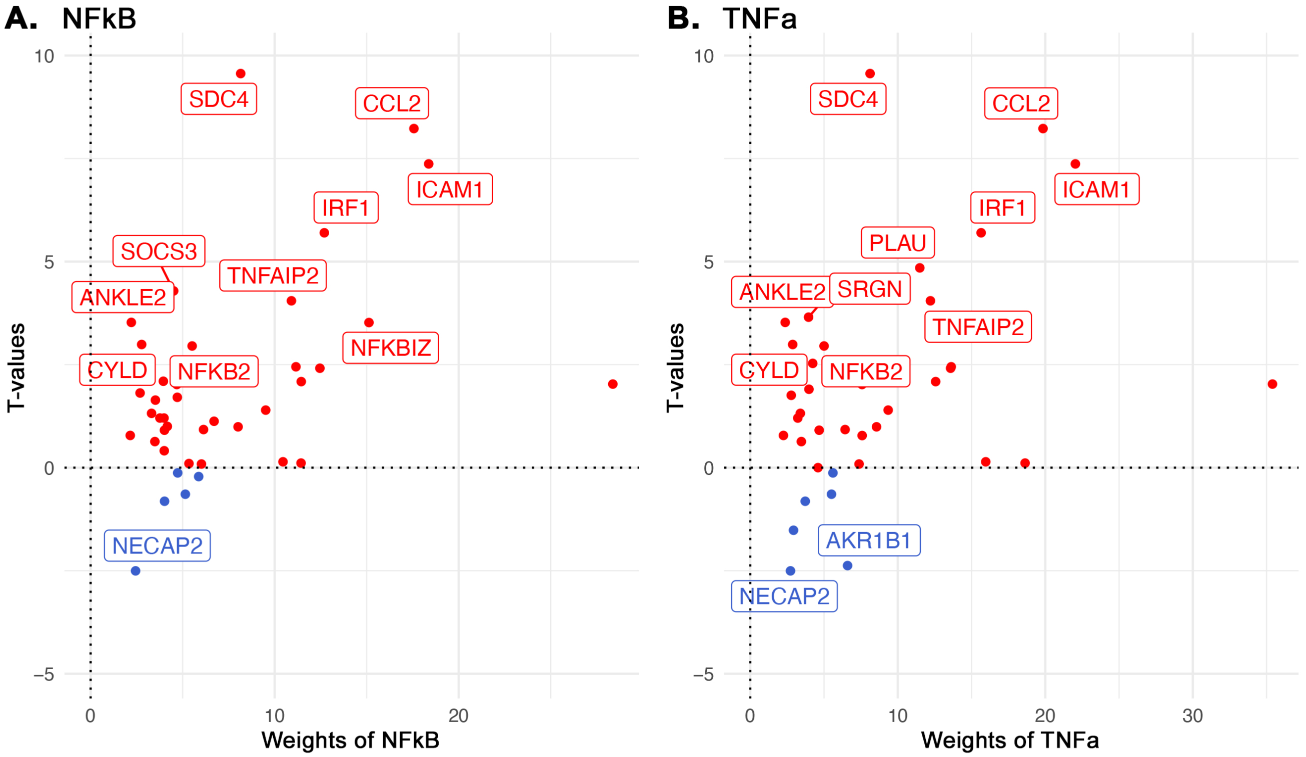


**Supplementary Figure 6.** Scatterplots of (A) NFkB and (B) TNFa pathway-responsive genes 4-hours following aerobic exercise. Pathway activity scores were calculated using PROGENy showing the association between pathway-responsive gene expression and the predicted activity of the signalling pathways. Red points indicate genes contributing positively to the pathway score, while blue points indicate genes contributing negatively to the pathway.

#### 1.2 Supplementary Tables

**Supplementary Table 1.** Canonical exercise sensor proteins.

| Sensors or receptors | Function | Protein | Gene | Targets |
|---|---|---|---|---|
| *Intracellular sensor proteins* | | | | |
| AMPK | Intracellular sensor of the [ATP]/[ADP] ratio | AMPK α-subunit 1 | *PRKAA1* | ☒ |
| | | AMPK α-subunit 2 | *PRKAA2* | ☒ |
| | | AMPK β-subunit 1 | *PRKAB1* | ☒ |
| | | AMPK β-subunit 2 | *PRKAB2* | ☒ |
| | | AMPK γ-subunit 1 | *PRKAG1* | ☒ |
| | | AMPK γ-subunit 2 | *PRKAG2* | ☒ |
| | | AMPK γ-subunit 3 | *PRKAG3* | ☒ |
| Ca2+, CaMKs | Sensor of changes in [$Ca^{2+}$] flux in the sarcoplasmic reticulum (required for muscle contraction) | Calmodulin-1 | *CALM1* | ☒ |
| | | Calmodulin-2 | *CALM2* | ☒ |
| | | Calmodulin-3 | *CALM3* | ☒ |
| | | Calcineurin subunit B type 1 | *PPP3R1* | ☒ |
| | | Calcineurin subunit B type 2 | *PPP3R2* | ☐* |
| cAMP | Sensor of cyclic AMP (cAMP) | Protein kinase A | *PRKACA* | ☒ |
| MAP3Ks | Comprise a large group of kinases that act as sensors and integrators of diverse signals, including mechanical stress, calcium flux, metabolic stress, oxidative stress, and cytokines | MEKK1 | *MAP3K1* | ☒ |
| | | MEKK2 | *MAP3K2* | ☒ |
| | | MEKK3 | *MAP3K3* | ☒ |
| | | MEKK4 | *MAP3K4* | ☒ |
| | | ASK1 | *MAP3K5* | ☒ |
| | | ASK2 | *MAP3K6* | ☒ |
| | | TAK1 | *MAP3K7* | ☒ |
| | | Tpl-2 | *MAP3K8* | ☒ |
| | | MLK1 | *MAP3K9* | ☐* |
| | | MLK2 | *MAP3K10* | ☒ |
| | | MLK3 | *MAP3K11* | ☒ |
| | | DLK | *MAP3K12* | ☒ |
| | | TAO1/2 | *TAOK1* | ☒ |
| Mechano-sensors | Sense changes in mechanical tension across the muscle | FAK | *PTK2* | ☒ |
| | | Yap | *YAP1* | ☒ |
| | | Diacylglycerol kinase zeta | *DGKZ* | ☒ |

| | | | | |
|---|---|---|---|---|
| | | Titin | *TTN* | ☒ |
| | | Integrin alpha 7/beta 1 | *ITGA7* | ☒ |
| | | BAG family molecular chaperone regulator 3 | *BAG3* | ☒ |
| | | Filamin-C | *FLNC* | ☒ |
| Oxidative stress | Sensor of reactive oxygen species | Inhibitor of Nrf2 | *KEAP1* | ☒ |
| Prolyl hydroxylases | Enzymes that sense intracellular $PO_2$ | PHD1 | *EGLN1* | ☒ |
| | | PHD2 | *EGLN2* | ☒ |
| | | PHD3 | *EGLN3* | ☒ |
| | | FIH1 | *HIF1AN* | ☒ |
| Sirtuins | Intracellular sensor of the [NAD+]/[NADH] ratio | Cytosolic | *SIRT1* | ☒ |
| | | Mitochondrial | *SIRT3* | ☒ |
| *Hormonal receptors* | | | | |
| Adiponectin | Sensor of adiponectin | Adiponectin receptor protein 1 | *ADIPOR1* | ☒ |
| | | Adiponectin receptor protein 2 | *ADIPOR2* | ☒ |
| Androgen receptors | Sensor of androgenic hormones | Androgen receptor | *AR* | ☒ |
| β-adrenergic receptors | Sense and transduce signals encoded by catecholamine hormones and neurotransmitters | β1 adrenergic receptor | *ADRB1* | ☒ |
| | | β2 adrenergic receptor | *ADRB2* | ☒ |
| | | β3 adrenergic receptor | *ADRB3* | ☐* |
| Growth hormone receptor | Sensor of circulating growth hormone | Growth hormone receptor | *GHR* | ☒ |
| Insulin | Sensor of plasma insulin | Insulin receptor | *INSR* | ☒ |
| Insulin-like growth factor 1 receptor | Activated by IGF-1 in circulation and/or mechanogrowth factor acting in autocrine/paracrine | IGF-1 receptor | *IGF1R* | ☒ |
| Myostatin | Sensor of myostatin and activins | Activin receptor type-2A | *ACVR2A* | ☒ |
| | | Activin receptor type-2B | *ACVR2B* | ☒ |
| TFG-β | Sensor of TGF-β | TGF-β receptor type 1 | *TGFBR1* | ☒ |
| | | TGF-β receptor type 2 | *TGFBR2* | ☒ |

*Gene was not included in the list of perturbation targets provided to standard CARNIVAL because it was absent from the OmniPath prior knowledge network.

**Supplementary Table 2.** Repeatability analysis of standard CARNIVAL network inference. Eight iterations of network inference were performed – four using the aerobic exercise data and four using the resistance exercise data – each with *RPS6KB1*-specific transcription factors as terminal nodes. The perturbation targets included the canonical exercise sensors (n=50), with beta weight set to 0.2. The table reports the number of inferred node-to-node interactions and individual nodes inferred in each replicate run.

| Run | Solution pool (n) | Interactions (n) | Nodes (n) |
|---|---|---|---|
| | | *Aerobic exercise* | |
| 1 | 100 | 26 | 26 |
| 2 | 100 | 27 | 26 |
| 3 | 100 | 25 | 24 |
| 4 | 100 | 26 | 25 |
| | | *Resistance exercise* | |
| 5 | 100 | 51 | 37 |
| 6 | 100 | 51 | 36 |
| 7 | 100 | 51 | 37 |
| 8 | 100 | 51 | 37 |

**Supplementary Table 3.** Repeatability analysis of the network interactions of the standard CARNIVAL network inference. Eight iterations of network inferences were performed – four using the aerobic exercise data and four using the resistance exercise data – using the *RPS6KB1*-specific transcription factors as terminal nodes. The perturbation targets included the canonical exercise sensors (n=50), with beta weight set to 0.2. The table compares interactions across the inferred networks. Interaction details include the number of discrepancies between networks (i.e., node interactions not shared), the percentage of preserved network interactions, discrepancies in edge weights, the percentage of edge weight differences, the absolute difference in edge weight if differences exist reported as an average or median, the number of positive edge weight discrepancies with average difference, and the number of negative edge weight discrepancies with average differences.

| | Interactions | | | | | | |
|---|---|---|---|---|---|---|---|
| | **Discrepant interactions (n)** | **Preserved interactions** | **Discrepant edge weights (n)** | **Preserved edge weights** | **Average absolute difference (median)** | **Positive edge discrepancy (n, average)^** | **Negative edge discrepancy (n, average)^** |
| | | | | *Aerobic exercise* | | | |
| Run 1 vs 2 | 1 | 96% | 12 | 56% | 17.8 (11.5) | 4 (13.3) | 8 (-20.0) |
| Run 1 vs 3 | 1 | 96% | 11 | 59% | 23.3 (22.0) | 4 (12.3) | 7 (-29.6) |
| Run 1 vs 4 | 0 | 100% | 10 | 63% | 18.3 (10.5) | 4 (9.5) | 6 (-24.2) |
| Run 2 vs 3 | 2 | 93% | 12 | 56% | 19.6 (13.0) | 5 (18.4) | 7 (-20.4) |
| Run 2 vs 4 | 1 | 96% | 12 | 56% | 16.3 (9.0) | 8 (12.3) | 4 (-24.5) |
| Run 3 vs 4 | 1 | 96% | 11 | 59% | 15.2 (17.0) | 5 (21.8) | 6 (-9.7) |
| | | | | *Resistance exercise* | | | |
| Run 5 vs 6 | 2 | 96% | 36 | 31% | 10.9 (4.0) | 16 (12.6) | 20 (-9.5) |
| Run 5 vs 7 | 0 | 100% | 29 | 44% | 3.8 (2.0) | 17 (3.5) | 12 (-4.3) |
| Run 5 vs 8 | 0 | 100% | 30 | 42% | 3.2 (2.0) | 14 (3.1) | 16 (-3.3) |
| Run 6 vs 7 | 2 | 96% | 40 | 23% | 9.7 (2.5) | 19 (10.0) | 21 (-9.3) |
| Run 6 vs 8 | 2 | 96% | 41 | 21% | 9.2 (2.0) | 22 (8.1) | 19 (-10.5) |
| Run 7 vs 8 | 0 | 100% | 32 | 38% | 3.1 (1.0) | 12 (3.4) | 20 (-2.9) |

^Positive and negative edge discrepancies indicate increased or reduced subnetworks, respectively, in the comparison run containing the specific interaction. Both the number of discrepancies, and their average difference are reported.

**Supplementary Table 4.** Repeatability analysis of the node attributes of the standard CARNIVAL network inference. Eight iterations of network inferences were performed – four using the aerobic exercise data and four using the resistance exercise data – using the *RPS6KB1*-specific transcription factors as terminal nodes. The perturbation targets included the canonical exercise sensors (n=50), with beta weight set to 0.2. The table compares node attributes across the inferred networks. Node attribute details include the number of discrepancies between networks (i.e., nodes not shared), the percentage of preserved nodes, discrepancies in node average activities, the percentage of average weight discrepancies, the absolute difference in average activity if differences exist reported as an average or median, the number of positive activity discrepancies with average difference, and the number of negative activity discrepancies with average differences.

| | Node attributes | | | | | | |
|---|---|---|---|---|---|---|---|
| | **Discrepant nodes (n)** | **Preserved interactions** | **Discrepant average activity (n)** | **Preserved edge weights** | **Average absolute difference (median)** | **Positive activity discrepancy (n, average)^** | **Negative activity discrepancy (n, average)^** |
| | | | *Aerobic exercise* | | | | |
| Run 1 vs 2 | 1 | 96% | 10 | 62% | 21.7 (16.0) | 2 (26.0) | 8 (-20.6) |
| Run 1 vs 3 | 1 | 96% | 9 | 65% | 23.1 (18.0) | 4 (11.0) | 5 (-32.8) |
| Run 1 vs 4 | 0 | 100% | 8 | 69% | 22.4 (11.5) | 2 (23.5) | 6 (-22.0) |
| Run 2 vs 3 | 2 | 92% | 10 | 62% | 19.0 (12.0) | 5 (22.8) | 5 (-24.2) |
| Run 2 vs 4 | 1 | 96% | 10 | 62% | 20.4 (8.5) | 8 (14.5) | 2 (-44.0) |
| Run 3 vs 4 | 1 | 96% | 9 | 65% | 13.7 (13.0) | 5 (15.8) | 4 (-11.0) |
| | | | *Resistance exercise* | | | | |
| Run 5 vs 6 | 1 | 97% | 19 | 49% | 11.5 (3.0) | 11 (9.5) | 8 (-14.4) |
| Run 5 vs 7 | 0 | 100% | 17 | 54% | 1.5 (1.0) | 9 (1.2) | 8 (-1.8) |
| Run 5 vs 8 | 0 | 100% | 15 | 59% | 1.6 (1.0) | 6 (1.33) | 9 (-1.8) |
| Run 6 vs 7 | 1 | 97% | 22 | 41% | 9.8 (2.0) | 9 (12.4) | 13 (-8.0) |
| Run 6 vs 8 | 1 | 97% | 21 | 43% | 10.1 (2.0) | 8 (13.5) | 13 (-8.1) |
| Run 7 vs 8 | 0 | 100% | 14 | 62% | 1.5 (1.0) | 5 (1.6) | 9 (-1.4) |

^Positive and negative activity discrepancies indicate increased or reduced average activity of the subnetwork node, respectively, in the comparison run containing the node. Both the number of discrepancies, and their average difference are reported.

**Supplementary Table 5.** Transcription factors controlled by upstream canonical exercise sensors. Interpretation of canonical exercise sensor regulation of downstream transcription factors (TFs) following standard CARNIVAL network inference from aerobic or resistance exercise data. For each inferred exercise sensor, the regulated downstream TFs are listed. For each TF, the minimum edge weight and average node activity (avgAct) along the signaling pathway are recorded. If a node lacks consistent stimulatory or inhibitory activity, the non-zero activity is reported (i.e., 1 – zeroAct). When multiple pathways exist between a sensor and a TF, the pathway with the highest minimum edge weight and avgAct is reported, as it is more likely to represent the biologically relevant route in the inferred network.

| **Exercise sensor** | **Regulated transcription factor** | **Minimum edge weight** | **Minimum avgAct (non-zeroAct)** |
|---|---|---|---|
| | *Aerobic exercise* | | |
| *ACVR2B* | *CTCF*<br>*FOXA1*<br>*MEF2A* | 58.2<br>58.2<br>58.2 | 31.0<br>31.0<br>31.0 |
| *EGLN1* | *HIF1A* | 60.4 | -52.3 |
| *EGLN2* | *HIF1A* | 51.5 | -30.9 |
| *EGLN3* | *HIF1A* | 49.7 | -10.5 (22.6) |
| *HIF1AN* | *HIF1A* | 47.1 | -43.9 |
| *INSR* | *CTCF*<br>*ELF1*<br>*E2F2*<br>*E2F1*<br>*FOXA1*<br>*MEF2A*<br>*MYC*<br>*NRF1* | 66.4<br>66.4<br>66.4<br>66.4<br>66.4<br>66.4<br>66.4<br>66.4 | 66.4<br>66.4<br>66.4<br>66.4<br>66.4<br>66.4<br>66.4<br>66.4 |
| *MAP3K7* | | | |
| *PRKAA1* | *SNAI1* | 100 | -100 |
| *PRKACA* | *CTCF*<br>*ELF1*<br>*E2F2*<br>*E2F1*<br>*FOXA1*<br>*MEF2A*<br>*MYC*<br>*NRF1* | 33.6<br>33.6<br>33.6<br>33.6<br>33.6<br>33.6<br>33.6<br>33.6 | 33.6<br>33.6<br>33.6<br>33.6<br>33.6<br>33.6<br>33.6<br>33.6 |
| *SIRT1* | *CTCF*<br>*E2F1*<br>*FOXA1*<br>*MEF2A* | 43.9<br>1.0<br>43.9<br>43.9 | 24.8<br>24.8<br>24.8<br>24.8 |

| | | | |
|---|---|---|---|
| | *NRF1* | 43.9 | 24.8 |
| *TGFBR1* | *CTCF*<br>*FOXA1*<br>*MEF2A* | 40.8<br>40.8<br>40.8 | 23.1<br>23.1<br>23.1 |
| *Resistance exercise* | | | |
| *ACVR2B* | *CEBPA* | 5.1 | -7.5 |
| *AR* | *CEBPA*<br>*ELF1*<br>*ESR1*<br>*E2F1*<br>*NRF1*<br>*SNAI1* | 2.7<br>22.8<br>27.8<br>25.0<br>22.8<br>22.8 | -1.8<br>14.6 (40.9)<br>14.6 (40.9)<br>14.6 (40.9)<br>14.6 (40.9)<br>30.6 |
| *INSR* | *ELF1*<br>*ESR1*<br>*NRF1*<br>*SNAI1* | 39.7<br>39.7<br>39.7<br>30.6 | 52.4<br>52.4<br>52.4<br>30.6 |
| *MAP3K1* | *CEBPA*<br>*ELF1*<br>*E2F1*<br>*ESR1*<br>*NRF1*<br>*SNAI1* | 21.2*<br>14.2*<br>14.2*<br>14.2*<br>14.2*<br>14.2* | -29.0<br>-29.0<br>-29.0<br>-29.0<br>-29.0<br>-29.0 |
| *MAP3K2* | *CEBPA*<br>*ELF1*<br>*E2F1*<br>*ESR1*<br>*NRF1*<br>*SNAI1* | 16.2<br>14.2<br>14.2<br>14.2<br>14.2<br>14.2 | -15.7<br>-15.7<br>-15.7<br>-15.7<br>-15.7<br>-15.7 |
| *MAP3K7* | *CEBPA* | 4.1 | -8.5 |
| *MAP3K8* | *CEBPA*<br>*ELF1*<br>*E2F1*<br>*ESR1*<br>*NRF1*<br>*SNAI1* | 25.6<br>14.2<br>14.2<br>14.2<br>14.2<br>14.2 | -31.3<br>-31.3<br>-31.3<br>-31.3<br>-31.3<br>-30.6 |
| *PRKAA1* | *E2F2*<br>*HIF1A*<br>*SNAI1* | 89.9<br>89.9<br>89.9 | -83.8 (96.1)<br>-83.8 (96.1)<br>-83.8 (96.1) |
| *PRKACA* | *ELF1*<br>*ESR1*<br>*E2F1*<br>*E2F2*<br>*HIF1A* | 47.6<br>73.9<br>27.0<br>42.2<br>42.2 | 47.6<br>69.8 (78.0)<br>47.6<br>47.4<br>47.4 |

| | | | |
|---|---|---|---|
| | *NRF1*<br>*SNAI1* | 47.6<br>13.3 | 47.6<br>30.6 |
| *SIRT1* | *E2F1* | 75.0 | -75.0 |
| *TGFBR1* | *CTCF*<br>*MEF2A* | 100.0<br>100.0 | 100.0<br>100.0 |

**Supplementary Table 6.** Combined control of transcription factors by exercise sensors. Interpretation of transcription factor (TF) regulation by upstream canonical exercise sensors following standard CARNIVAL network inference from aerobic or resistance exercise data. For each TF, the upstream exercise sensors that regulate the TF are listed. For each sensor, the minimum edge weight and average node activity (avgAct) along the signaling pathway to the TF are documented. If a node lacked consistent stimulatory or inhibitory activity, the non-zero activity is reported (i.e., 1 – zeroAct) to indicate inconsistency in activity. When multiple pathways exist between a sensor and a TF, the pathway with the highest minimum edge weight and avgAct is reported, as it is more likely to represent the biologically relevant route in the inferred network.

| **Transcription factor** | **Regulatory exercise sensors (minimum edge weight, minimum actAvg [non-zeroAct])** | |
|---|---|---|
| | *Aerobic exercise* | |
| *CTCF* | *ACVR2B* (58.2, 31.0)<br>*INSR* (66.4, 66.4)<br>*PRKACA* (33.6, 33.6) | *SIRT* (43.9, 24.8)<br>*TGFBR1* (40.8, 23.1) |
| *ELF1* | *INSR* (66.4, 66.4) | *PRKACA* (33.6, 33.6) |
| *E2F1* | *INSR* (66.4, 66.4)<br>*PRKACA* (33.6, 33.6) | *SIRT1* (1.0, 24.8) |
| *E2F2* | *INSR* (66.4, 66.4) | *PRKACA* (33.6, 33.6) |
| *FOXA1* | *ACVR2B* (58.2, 31.0)<br>*INSR* (66.4, 66.4)<br>*PRKACA* (33.6, 33.6) | *SIRT1* (43.9, 24.8)<br>*TGFBR1* (40.8, 23.1) |
| *HIF1A* | *EGLN1* (60.4, -52.3)<br>*EGLN2* (51.5, -30.9) | *EGLN3* (49.7, -10.5 [22.6])<br>*HIF1AN* (47.1, -43.9) |
| *MEF2A* | *ACVR2B* (58.2, 31.0)<br>*INSR* (66.4, 66.4)<br>*PRKACA* (33.6, 33.6) | *SIRT* (43.9, 24.8)<br>*TGFBR1* (40.8, 23.1) |
| *MYC* | *INSR* (66.4, 66.4) | *PRKACA* (33.6, 33.6) |
| *NRF1* | *INSR* (66.4, 66.4)<br>*PRKACA* (33.6, 33.6) | *SIRT* (43.9, 24.8) |
| *SNAI1* | *PRKAA1* (100, -100) | |
| | *Resistance exercise* | |
| *CEBPA* | *ACVR2B* (5.1, -7.5)<br>*MAP3K1* (21.2, -29.0)<br>*MAP3K2* (16.2, -15.7) | *MAP3K7* (4.1, -8.5)<br>*MAP3K8* (25.6, -31.3) |
| *CTCF* | *TGFBR1* (100, 100) | |
| *ELF1* | *AR* (22.8, 14.6 [40.9])<br>*INSR* (39.7, 52.4)<br>*MAP3K1* (14.2, -29.0) | *MAP3K2* (14.2, -15.7)<br>*MAP3K8* (14.2, -31.3)<br>*PRKACA* (47.6, 47.6) |
| *ESR1* | *AR* (27.8, 14.6 [40.9]) | *MAP3K2* (14.2, -15.7) |

| | | |
|---|---|---|
| | *INSR* (39.7, 52.4)<br>*MAP3K1* (14.2, -29.0) | *MAP3K8* (14.2, -31.3)<br>*PRKACA* (73.9, 69.8 [78.0]) |
| *E2F1* | *AR* (25.0, 14.6 [40.9])<br>*MAP3K1* (14.2, -29.0)<br>*MAP3K2* (14.2, -31.3) | *MAP3K8* (14.2, -31.3)<br>*PRKACA* (27.0, 47.6)<br>*SIRT1* (75.0, -75.0) |
| *E2F2* | *PRKAA1* (89.9, -83.8 [96.1]) | *PRKACA* (42.2, 47.6) |
| *HIF1A* | *PRKAA1* (89.9, -83.8 [96.1]) | *PRKACA* (42.2, 47.6) |
| *MEF2A* | *TGFBR1* (100, 100) | |
| *NRF1* | *AR* (22.8, 14.6 [40.9])<br>*INSR* (39.7, 52.4)<br>*MAP3K1* (14.2, -29.0) | *MAP3K2* (14.2, -15.7)<br>*MAP3K8* (14.2, -31.3)<br>*PRKACA* (47.6, 47.6) |
| *SNAI1* | *AR* (22.8, 30.6)<br>*INSR* (30.6, 30.6)<br>*MAP3K1* (14.2, -29.0)<br>*MAP3K2* (14.2, -15.7) | *MAP3K8* (14.2, -30.6)<br>*PRKAA1* (89.9, -83.8 [96.1])<br>*PRKACA* (13.3, 30.6) |
